# Human iPSC-derived cerebral organoids model features of Leigh Syndrome and reveal abnormal corticogenesis

**DOI:** 10.1101/2020.04.21.054361

**Authors:** Alejandra I. Romero-Morales, Gabriella L. Robertson, Anuj Rastogi, Megan L. Rasmussen, Hoor Temuri, Gregory Scott McElroy, Ram Prosad Chakrabarty, Lawrence Hsu, Paula M. Almonacid, Bryan A. Millis, Navdeep S. Chandel, Jean-Philippe Cartailler, Vivian Gama

## Abstract

Leigh syndrome (LS) is a rare, inherited neurometabolic disorder that presents with bilateral brain lesions, caused by defects in the mitochondrial respiratory chain and associated nuclear-encoded proteins. We generated iPSCs from three patient-derived LS fibroblast lines and identified, by whole-exome and mitochondrial sequencing, unreported mutations in pyruvate dehydrogenase (GM0372, PDH; GM13411, MT-ATP6/PDH) and dihydrolipoyl dehydrogenase (GM01503, DLD). LS-derived iPSC lines were viable and generally capable of differentiating into key progenitor populations, but we identified several abnormalities in three-dimensional differentiation models of brain development. LS-derived cerebral organoids showed defects in neural epithelial bud generation, size, and cortical architecture at 100 days. The double mutant MT-ATP6/PDH line produced organoid neural progenitor cells with abnormal mitochondrial morphology characterized by fragmentation and disorganization and showed an increased generation of astrocytes. These studies aim to provide a comprehensive phenotypic characterization of available patient-derived cell lines that can be used as LS model systems.

## Introduction

Leigh syndrome (LS), or sub-acute necrotizing encephalomyelopathy, is an inherited neurometabolic disorder that affects the central nervous system (CNS) (Baertling et al., 2014; Gerards et al., 2016; Leigh, 1951; Sorbi and Blass, 1982). LS is a rare, progressive, early-onset disease with a prevalence of 1 in 40,000 live births (Lake et al., 2016). The pathologic features of LS are focal, bilateral lesions in one or more areas of the CNS, including the brainstem, thalamus, basal ganglia, cerebellum, and spinal cord. The most common underlying cause is defective oxidative phosphorylation (OXPHOS), due to mutations in genes encoding complexes of the mitochondrial respiratory chain (Baertling et al., 2014; Lake et al., 2015, 2016). The LS phenotype caused by mitochondrial-encoded gene mutations can be rescued by mitochondrial replacement, highlighting the importance of this organelle in LS disease progression (Ma et al., 2015).

The availability of animal models and brain tissue from biopsies has provided critical insight into this disease. However, our understanding of the etiology and pathology of complex neurological diseases like LS would benefit from human-derived platforms such as induced pluripotent stem cell-derived models (Inak et al., 2021; Lorenz et al., 2017; Quadrato et al., 2016). The ability to reprogram somatic cells into induced pluripotent stem cells (iPSCs), followed by differentiation into specific lineages has become a useful tool for complex disease modeling (Kelava and Lancaster, 2016; Di Lullo and Kriegstein, 2017; Pașca, 2018; Quadrato et al., 2016). In the context of LS, iPSCs have been successfully generated from patients with mutations in mitochondrially encoded ATP Synthase Membrane Subunit 6 (MT-ATP6) (Galera-Monge et al., 2016; Grace et al., 2019; Lorenz et al., 2017; Ma et al., 2015), mitochondrially encoded NADH:Ubiquinone Oxidoreductase Core Subunit 3 (MT-ND3) subunit (Hattori et al., 2016) and the nuclear-encoded gene Surfeit locus protein 1 (SURF1) (Inak et al., 2019, 2021). These iPSC-model systems have been proposed for drug discovery (Inak et al., 2017; Lorenz et al., 2017) as well as testing platforms for potential metabolic rescue treatments (Ma et al., 2015).

Many studies have used LS patient fibroblasts commercially available at the Coriell Institute (Galera-Monge et al., 2016; Hinman et al., 1989; Huh et al., 1990; Iyer et al., 2012; Johnson et al., 2019; Ma et al., 2015; Sorbi and Blass, 1982; Vo et al., 2007; Zheng et al., 2016a). Here we report our findings on the genomic and phenotypic characterization of iPSCs derived from these three LS fibroblast lines. Whole exome and mitochondrial sequencing revealed previously unidentified mutations in these patient-derived cell lines. Three-dimensional differentiation of these LS mutant iPSCs into neural rosettes and cerebral organoids resulted in severe abnormalities. LS-iPSC-derived 100-day cerebral organoids showed decreased size as well as defects in the generation of neural epithelial buds. Corticogenesis was impaired in all LS mutant cell lines. MT-ATP6/PDH showed a decrease in gene expression in neural progenitor and cortical plate markers at day 30, while all three LS lines showed a reduction of the cortical layer markers CTIP2, TBR1, SATB2, and BRN2 at day 100. These results point to aberrant corticogenesis as a driver of LS pathogenesis and demonstrate the utility of iPSC-derived systems to recapitulate CNS phenotypes and to test potential strategies to restore neurogenesis in LS.

## Results

### Genomic characterization of Leigh syndrome fibroblasts

Due to the limited genomic information available for the three cell lines, we performed whole-exome sequencing (WES) and mitochondrial sequencing of the fibroblasts before reprogramming (Figure 1A-D and Supplemental Figure 1; data repository with the raw and analyzed WES and mitochondrial results can be found at https://www.ncbi.nlm.nih.gov/sra/PRJNA626388 & https://vandydata.github.io/Romero-Morales-Gama-Leigh-Syndrome-WES/). Comparison between the high impact, moderate impact, and all variants for identified INDELs and SNPs showed a significant overlap between the three cell lines (Supplemental Figure 1A & B). In-depth analysis of the top 15 high-impact SNPs (Supplemental Figure 1C) also confirmed an overlap between genotypes, with only three genes with confirmed SNPs related to neurological diseases (*FRG2C* for bipolar disorder and *CDC27* and *KIR2DL4* for white matter microstructure measurements) (Buniello et al., 2019).

**Figure 1.**
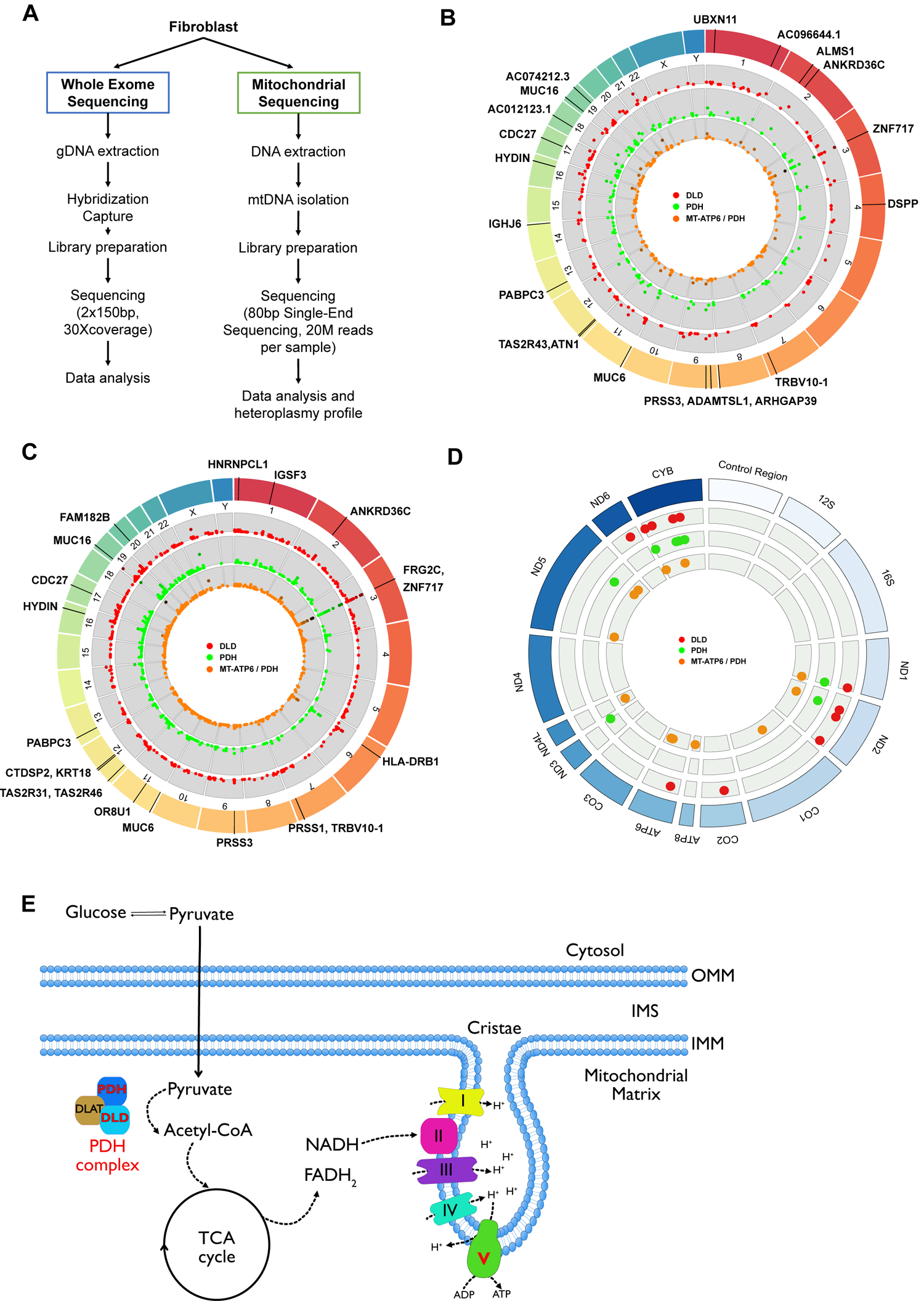
Whole exome sequencing identifies novel mutations in Leigh syndrome fibroblasts. A. Schematic of the WES and mitochondrial sequencing workflow. B. Representation of whole-genome sequencing data, highlighting the top 20 genes containing high impact indels. C. Representation of whole-genome sequencing data, highlighting the top 20 genes containing high impact SNPs (increased likelihood of disrupting protein function). D. Mitochondrial sequencing identifies novel mutations in LS fibroblasts. Representation of mitochondrial sequencing data, highlighting mitochondrial genes containing mutations (transitions, deletions, or transversions). Red dots: represent the DLD line. Green dots: represent the PDH line. Orange dots: represent the MT-ATP6/PDH line. E. Representation of the affected proteins in the LS mutant cell lines. PDH and DLD are part of the pyruvate dehydrogenase complex (PDHc). MT-ATP6 is a subunit of the ATP synthase, represented here as the electro transport chain complex V. PDH: Pyruvate dehydrogenase. DLD: Dihydrolipoyl dehydrogenase. MT-ATP6/PDH: Mitochondrially Encoded ATP Synthase Membrane Subunit 6/ Pyruvate dehydrogenase.

Targeted analysis of the genes associated with Leigh syndrome (Lake et al., 2016) revealed a loss of function insertion/deletion (INDEL) frameshift in pyruvate dehydrogenase complex (PDHc) E1 alpha 1 subunit or pyruvate dehydrogenase (PDHA1, c.79delC, p.Arg27fs) in the cell lines GM03672 and GM13411 (Figure 1E). A single-nucleotide polymorphism (SNP) in the PDHc E3 subunit or dihydrolipoyl dehydrogenase (DLD, c.100A>G, p.Thr34Ala) was identified in GM01503 (Figure 1E). In addition to being part of the PDHc, DLD is also a component of the α-ketoglutarate and branched-chain α-ketoacid dehydrogenase complexes (Craigen, 1996). Despite the lack of genomic data, dysfunction of the PDH complex was previously suggested in GM03672 and GM01503 as the main driver of the disease in these patients (Hinman et al., 1989; Huh et al., 1990; Sorbi and Blass, 1982) (Supplemental Table 1). To our knowledge, mutations in the nuclear genome of GM13411 have not been reported to date.

Mitochondrial sequencing identified several SNPs in all the cell lines (Figure 1D). A loss of function SNP in the MT-ATP6 gene was identified in the GM13411 line. This mutation was reported in the original clinical case (Pastores et al., 1994). The authors described the T to G transition at the 8993 position that results in the substitution of a highly conserved leucine residue for an arginine (L156R). MT-ATP6 is part of the F0 domain of ATP synthase that functions as a proton channel (Figure 1E). The L156R substitution prevents the induction of c-ring rotation (rotation of an oligomeric assembly of c-subunits of the ATP synthase (Kühlbrandt and Davies, 2016)), resulting in decreased ATP synthesis (Uittenbogaard et al., 2018). Heteroplasmy analysis of fibroblasts showed a 92% frequency of this mutation in the cell population, which is consistent with previous reports (Galera-Monge et al., 2016; Iyer et al., 2012; Pastores et al., 1994).

### Characterization of iPSCs derived from commercially available Leigh syndrome fibroblasts

Metabolic remodeling is a crucial step toward reprogramming somatic cells into iPSCs (Mathieu and Ruohola-Baker, 2017; Panopoulos et al., 2012; Rasmussen and Gama, 2020; Rastogi et al., 2019; Wu et al., 2016). Reprogramming of fibroblasts was performed as previously described (Takahashi et al., 2007) (Supplemental Figure 2A); we generated iPSC clones from a healthy age-matched control and the three LS cell lines. Pluripotency was evaluated using the microarray-based analysis PluriTest (Müller et al., 2011). All three LS cell lines showed a high pluripotency score and a low novelty score (Supplemental Figure 2B-C), congruent with the transcriptional profile of pluripotent stem cells. Moreover, all the reprogrammed cells expressed the pluripotent markers *NANOG* and *POU5F1* (*OCT4)* (Supplemental Figure 2D) and had a normal karyotype. The MT-ATP6/PDH cell line showed increased levels of NANOG (p<0.0001) compared to the control.

To assess the ability of the LS and control cell lines to differentiate into the three germ layers, we performed trilineage differentiation as previously described (Kuang et al., 2019; Roberts et al., 2019). Commitment to ectodermal fate was evaluated by immunofluorescence staining of the marker PAX6 (Figure 2A and B), as well as the mRNA expression of the genes *GATA3* and *PAX6* using RT-qPCR (Figure 2C). The endoderm lineage was assessed by protein expression of the marker SOX17 (Figure 2A and B) and by the expression of the genes *CDX2* and *SOX17* (Figure 2C). Finally, mesodermal lineage was confirmed by the staining of the protein markers Brachyury and CXCR4 (Figure 2A and B) and the expression of the genes *TBXT* and *NCAM* (Figure 2C). No significant differences were detected among the different lines when compared to control. Clinical data available from the patient with the PDH mutation reports elevated blood pyruvate levels (Schubert and Vilarinho, 2020; Sofou et al., 2014). High concentrations of pyruvate in the media have been shown to potentiate the differentiation of human embryonic stem cells (hESCs) into endodermal and mesodermal lineages in a lineage-specific fashion (Song et al., 2019), suggesting a differential capacity of these cells to preferentially commit into certain cell fates due to metabolic dysregulation.

**Figure 2.**
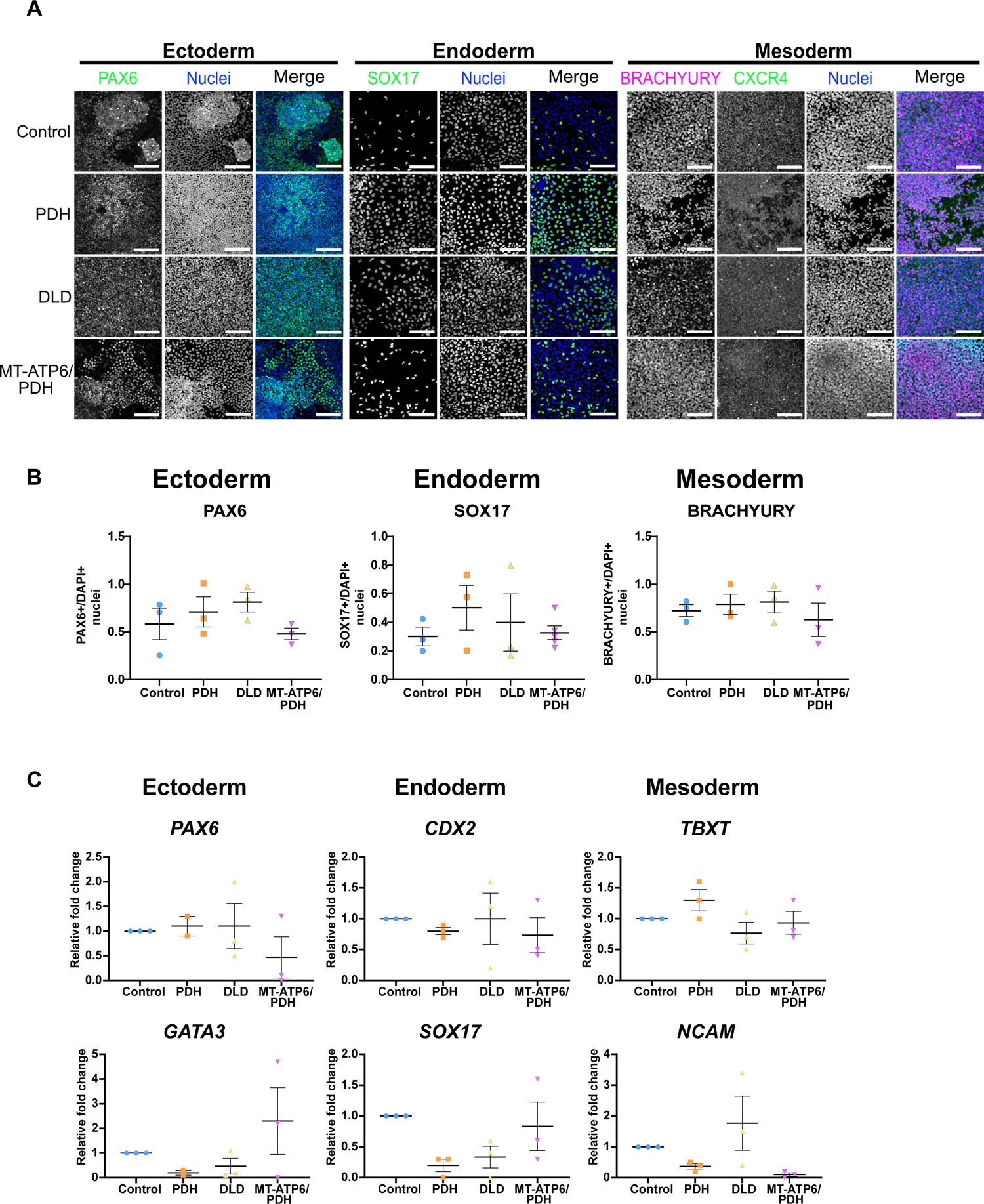
Induced pluripotent stem cells derived from Leigh syndrome patient fibroblasts are capable of differentiation into specific lineages. A. Representative images of the three embryonic lineages markers. Ectoderm marker Pax6, endoderm marker SOX17, and mesoderm markers Brachyury and CXCR4. Scale bar: 100μm. B. Immunofluorescence quantification for the aforementioned markers. Three independent differentiations were performed. Positive nuclei for the analyzed markers were divided by the total number of nuclei stained with DAPI. C. RT-qPCR analysis for the ectodermal genes *GATA3* and *PAX6*, ectodermal genes *CDX2* and *SOX17* the mesodermal genes *TBXT* and *NCAM*. Fold change normalized to GPI and GAPDH as housekeeping genes. Graphs represent mean ± SEM from at least three independent experiments.

### Two-dimensional neural differentiation is not significantly altered by Leigh syndrome-associated mutations

To determine if the LS mutations impact the commitment and development of the neural lineage, neural progenitor cells (NPCs) were generated by a dual SMAD inhibition protocol (Chambers et al., 2009) (Supplemental Figure 3A). NPCs expressed expected neural markers PAX6, NESTIN, and SOX2 (Figure 3A and Supplemental Figure 3B), with no significant differences between the LS cell lines and control (Figure 3B and Supplemental Figure 3B). The multipotent capacity of NPCs to generate neurons, astrocytes, and oligodendrocytes was evaluated using immunostaining and quantification of mRNA levels by RT-qPCR as previously reported (Chambers et al., 2009; TCW et al., 2017; Vescovi and Snyder, 1999). We identified a significant increase in the mean fluorescence intensity of the astrocyte marker S100β in the DLD mutant line (p=0.0185). These results suggest an increased capacity of these cells to commit to the astrocyte lineage. No other differences were observed at protein or mRNA level in the neuronal or oligodendrocyte progenitor lineage. (Figure 3A-C). Thus, LS NPCs are multipotent, capable of differentiation into the main cellular populations of the nervous system.

**Figure 3.**
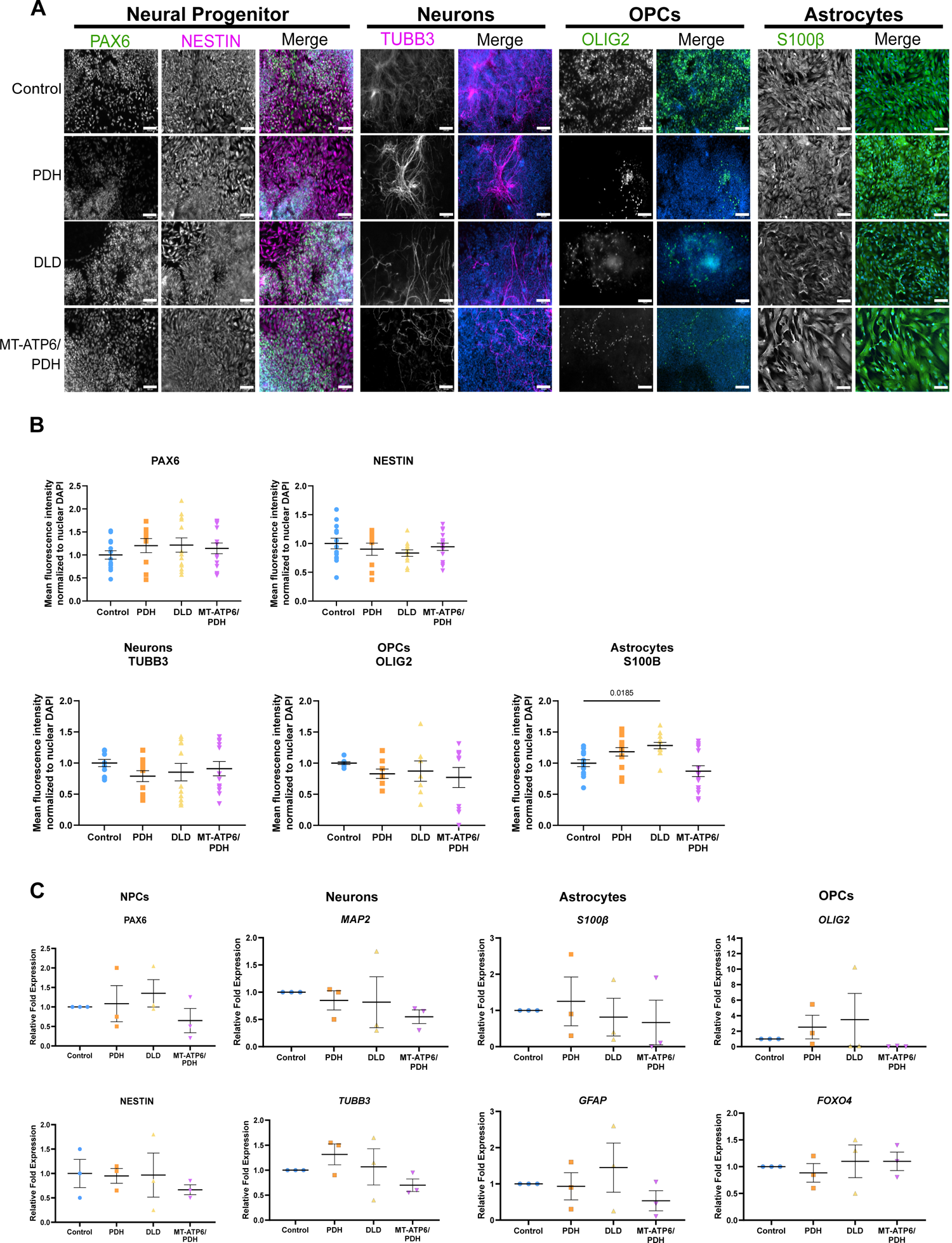
Leigh syndrome-derived NPCs are multipotent and do not show aberrant mitochondrial morphology compared to control. A. Representative images of the expression of neural multipotency markers. Neural progenitor cells stained by PAX6 and NESTIN, neurons marked with βIII-TUBULIN (TUBB3), oligodendrocyte progenitor cells (OPCs) were stained with Olig2, and astrocytes are marked with S100β. Merged panels show the color image of the grayscale lineage marker and the nuclear staining DAPI in blue. Scale bar: 100μm. B. Immunofluorescence quantification for the aforementioned markers. Three independent differentiations were performed. Mean fluorescence intensity for each marker was normalized to the nuclear DAPI intensity and the intensity values of control. C. RT-qPCR Analysis of the NPC markers *PAX6* and *NESTIN*, as well as the multipotency markers *MAP2* and *TUBB3* for neuronal lineage, *S100B* and *GFAP* for astrocytic lineage, and *OLIG2* and *FOXO4* for OPCs. Fold change normalized to GPI and GAPDH as house-keeping genes.

Neural cell death is a hallmark of LS; thus, we performed a cell viability assay to investigate the sensitivity of the LS-derived NPCs to different apoptotic stimuli (Supplemental Figure 3C). Treatment with DNA damaging agents, etoposide and neocarzinostatin, as well as the microtubule depolymerizing agent nocodazole, did not show increased sensitivity to cell death in LS NPCs compared to control. To evaluate the susceptibility of the NPCs to mitochondrial damage, we treated these cells with the mitochondrial oxidative phosphorylation uncoupler, CCCP. No difference was observed in the viability of the LS NPCs compared to control-treated with this mitochondrial toxicant.

### LS mutations cause morphological alterations in three-dimensional models of neurodevelopment

Previous studies using cells from LS patients carrying homozygous *SURF1* (c.769G>A and c.530T>G) and MT-ATP6 (m.9185T>C) mutations showed an abnormal generation of neural lineages (Lorenz et al., 2017) and impaired neurogenesis in cerebral organoids (Inak et al., 2019). Therefore, we investigated the effects of the PDH, DLD, and MT-ATP6/PDH LS mutations on neurogenesis using three-dimensional models of neural development as described previously (Lancaster and Knoblich, 2014; Romero-Morales et al., 2019). Most of these three-dimensional neural systems start with the formation of neuralized embryoid bodies (EBs). EBs are generated from a single cell suspension (Itskovitz-Eldor et al., 2000) plated in ultra-low attachment microwells with neural induction media (Figure 4A). EB diameter was measured on day 5 before plating on Matrigel-coated imaging plates. All three LS mutant lines showed increased diameter compared to control (Figure 4B, PDH vs control & DLD vs control: p<0.0001, MT-ATP6/PDH vs control: p=0.0209).

**Figure 4.**
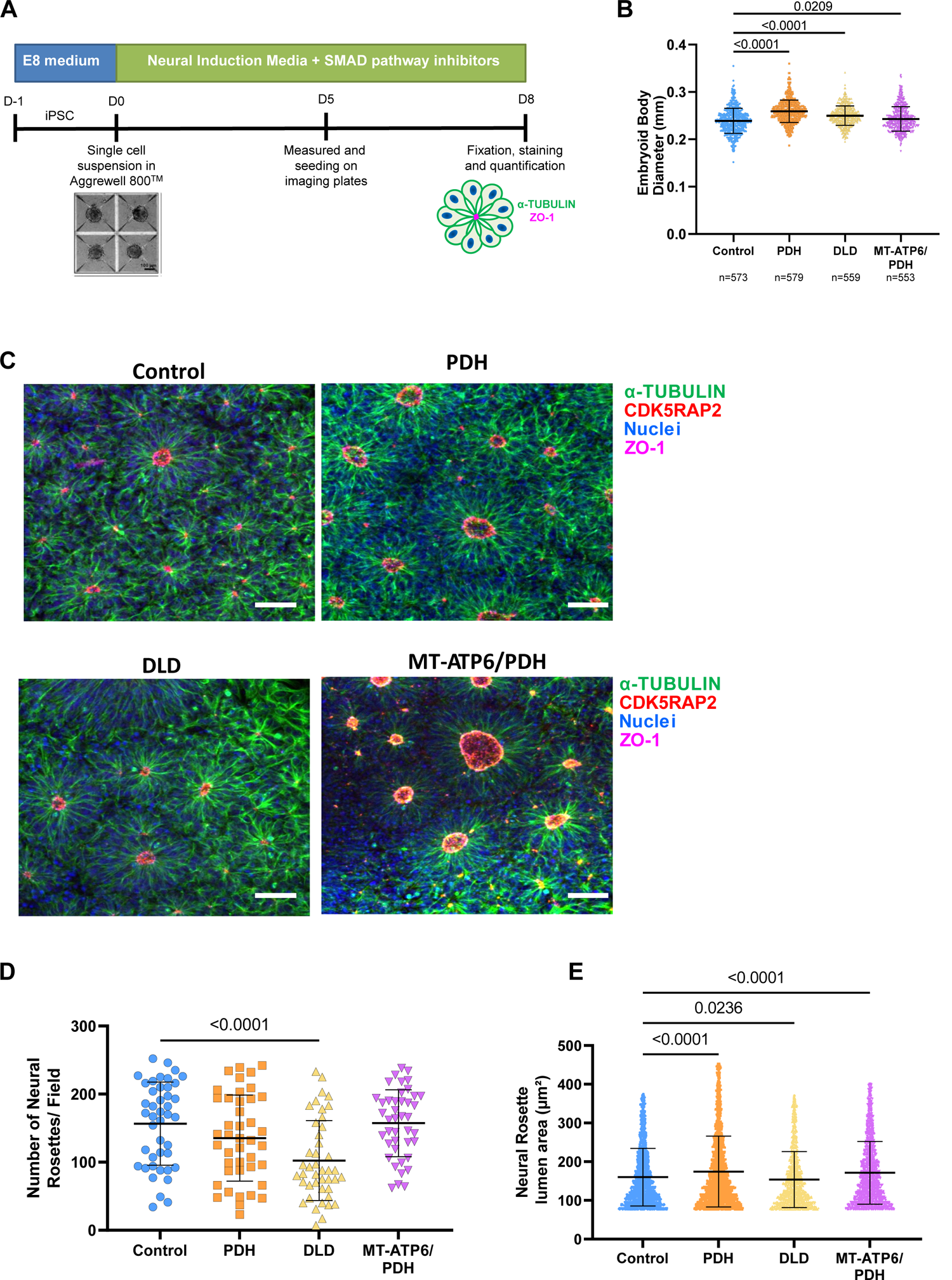
Three-dimensional differentiation reveals abnormalities during induction of neural rosettes in LS cell lines. A. Schematic of neural rosette generation protocol. B. Embryoid body diameter is increased in the three LS cell lines. C-E. Representative confocal images of neural rosettes (C) show decreased numbers of neural rosettes per field in the DLD mutant line (D). Quantification of the lumen area (μm^2^) indicates increased lumen area in the PDH and MT-ATP6/PDH mutant cell lines and a decreased lumen area in the DLD mutant line (E). Scale bar: 50μm.

To investigate the effects of LS-associated mutations in the early stages of CNS development, we generated neural rosettes using EBs grown in the presence of SMAD inhibitor media (Figure 4A) (Elkabetz et al., 2008; Zhang et al., 2001). These structures have previously been established to recapitulate the early neural tube formation stage of development (Elkabetz et al., 2008; Wilson and Stice, 2006). Neural rosettes were stained with the tight junction marker ZO-1 (Elkabetz et al., 2008; Hříbková et al., 2018) and the centrosomal marker CDK5RAP2 (Figure 4C). Quantification of the number of neural rosettes per field of view showed a reduction in these structures in the DLD mutant compared to controls (Figure 4D, p<0.001). Consistent with the overall increase in EB diameter (Figure 4B), lumen area quantification showed an increase in the PDH and MT-ATP6/PDH mutants, while the DLD mutant line showed a decrease in their area relative to controls (Figure 4E PDH: p<0.0001, DLD: p=0.0236 and MT-ATP6/PDH: p<0.0001). The neural rosettes obtained from all cell lines followed the expected morphological changes described previously (Hříbková et al., 2018). The polymerization of α-tubulin and the generation of the ZO-1 ring at the apical region of the rosettes are conserved in the LS mutants. Increased neural rosette lumen size has previously been associated with activation of the TGFβ pathway (Medelnik et al., 2018), Notch and sonic hedgehog (SHH) pathway, and inhibition of WNT i (Elkabetz et al., 2008). Large rosette formation is thought to be a consequence of coalescence or fusion of smaller rosettes (Fedorova et al., 2019) or apical domain opening and expansion rather than a process dependent on cell proliferation (Medelnik et al., 2018).

### Leigh syndrome-associated mutations disrupt corticogenesis in cerebral organoids

To investigate the effects of these mutations during corticogenesis, we generated cerebral organoids from LS iPSCs (Supplemental Figure 4A). Differences between the cell lines became apparent as early as the neuroepithelial bud expansion phase. After Matrigel embedding, the MT-ATP6/PDH mutant cell line showed poor budding with large areas of non-neuroepithelial cells (Supplemental Figure 4B). Defective organoid formation in this cell line was significantly higher than control and the other two LS cell lines (Supplemental Figure 4C). A previous report showed that when iPSCs generated from fibroblasts harboring the same T8993G mitochondrial mutation were differentiated into EBs, there was rapid regression and death after 7 days in suspension, while the monolayer culture did not show obvious deficits in cell growth (Grace et al., 2019). Given that the neuroectoderm expansion phase happens during days 7-10, the degeneration of the MT-ATP6/PDH organoids after embedding may be consistent with these reports. Higher metabolic requirements are associated with NPC proliferation and migration in three-dimensional scaffolds and development (Fang et al., 2020; Homem et al., 2015). As the PDH mutant line did not show this particular phenotype during the expansion of the neural epithelium, the presence of the additional mitochondrial mutation in the MT-ATP6/PDH line resulted in the reduction of organoid formation efficiency.

Cerebral organoid growth was tracked for 100 days (Supplemental Figure 4D). Growth differences in the average diameter of the organoids became significant at day 80 between the control and the PDH mutant line (p< 0.05). At days 90 and 100, organoids derived from all LS cell lines were significantly smaller than the controls (p< 0.05). Phenotypic information available about the patient from which the PDH mutant cell line was derived indicates that this particular patient presented with microcephaly. Hence, we were able to mimic an aspect of the patient phenotype *in vitro* using the cerebral organoid model system.

To assess the effect of the LS mutations on development during the first stages of neural development, we collect mRNA of day 30 organoids to evaluate the pattern of expression of the different NPC and cortical markers by RT-qPCR (Supplemental Figure 5A). NPC marker *SOX2* expression was reduced in all three LS mutants (PDH: p=0.0156, DLD p=0.0303, MT-ATP6/DPH: p<0.0001). Progenitor markers *NESTIN* and *PAX6* expression were also increased in PDH mutant organoids (p=0.0156 and p=0.0134, respectively). In the case of MT-ATP6/PDH organoids, there was a significant reduction in the expression of *PAX6* (p=0.0231) and a significant increase in the expression of the intermediate progenitor cell (IPC) marker *TBR2* (p=0.0224). The cortical plate marker *CTIP2* was found to be reduced in both DLD (p=0.0080) and MT-ATP6/PDH (p<0.0001); and the neuronal marker βIII-TUBULIN (*TUBB3*) was reduced in MT-ATP6/PDH (p=0.0302). No significant differences were noted in the expression of the glycoprotein *REELIN* or the cortical plate marker *TBR1* among the different genotypes.

The reduction in the expression of the neural progenitor markers *SOX2* and *PAX6* in the MT-ATP6/PDH concomitantly with the increase in *TBR2* may suggest a premature commitment into intermediate progenitor cells (IPCs) (Englund, 2005; Hutton and Pevny, 2011; Sansom et al., 2009). This premature differentiation into IPCs and the reduced expression of committed neuronal markers such as *CTIP2* and *TUBB3* may suggest an inability to acquire a neuronal fate in this particular genotype.

To assess if the differences observed at an mRNA level were maintained at the protein level, as well as to analyze the overall cytoarchitecture of the organoids at day 30 timepoint, brain organoids were sectioned and stained for the ventricular zone (VZ), subventricular zone (sVZ), and cortical plate (CP) markers (Supplemental Figure 5 B-E). Day 30 organoids were obtained from at least 3 independent batches of differentiation and representative images were obtained from at least 4 individual organoids per batch. Quantification of the immunofluorescence images revealed no significant differences among the number of NPC cells positive for the markers SOX2, PAX6, and NESTIN or IPC marker TBR2 (Supplemental Figure 5F). In agreement with the defective neuroepithelial expansion, the overall architecture in MT-ATP6/PDH organoids was compromised. Few ventricle-like structures were present, and the foci of PAX6+ cells were not organized in the expected radial pattern. Migration of early-born neurons *in vivo* depends on pioneer Cajal-Retzius neurons that are positive for the glycoprotein REELIN (Lancaster et al., 2017). Cells positive for this marker were identified in superficial regions of all organoids. Early-born neurons at this time point are expected to migrate into the CP that will later give rise to the deep cortical layers (Camp et al., 2015). CTIP2+ and TBR1+ cells were observed in all genotypes. The neuronal marker Microtubule-associated protein 2 (MAP2) was also present in all samples at similar levels to control (Supplemental Figure 5E and 5F). The outer radial glia marker Homeodomain-only protein (HOPX) expression was significantly reduced in the PDH (p<0.0001) and MT-ATP6/PDH mutant (p=0.0417). Metabolic stress has been correlated with reduced specification in organoids, especially in oRG and newborn neurons (Bhaduri et al., 2020). Hence, lower levels of HOPX positive cells in the cell lines harboring a PDH mutation may be associated with defects in cellular fate specification at this timepoint.

**Figure 5.**
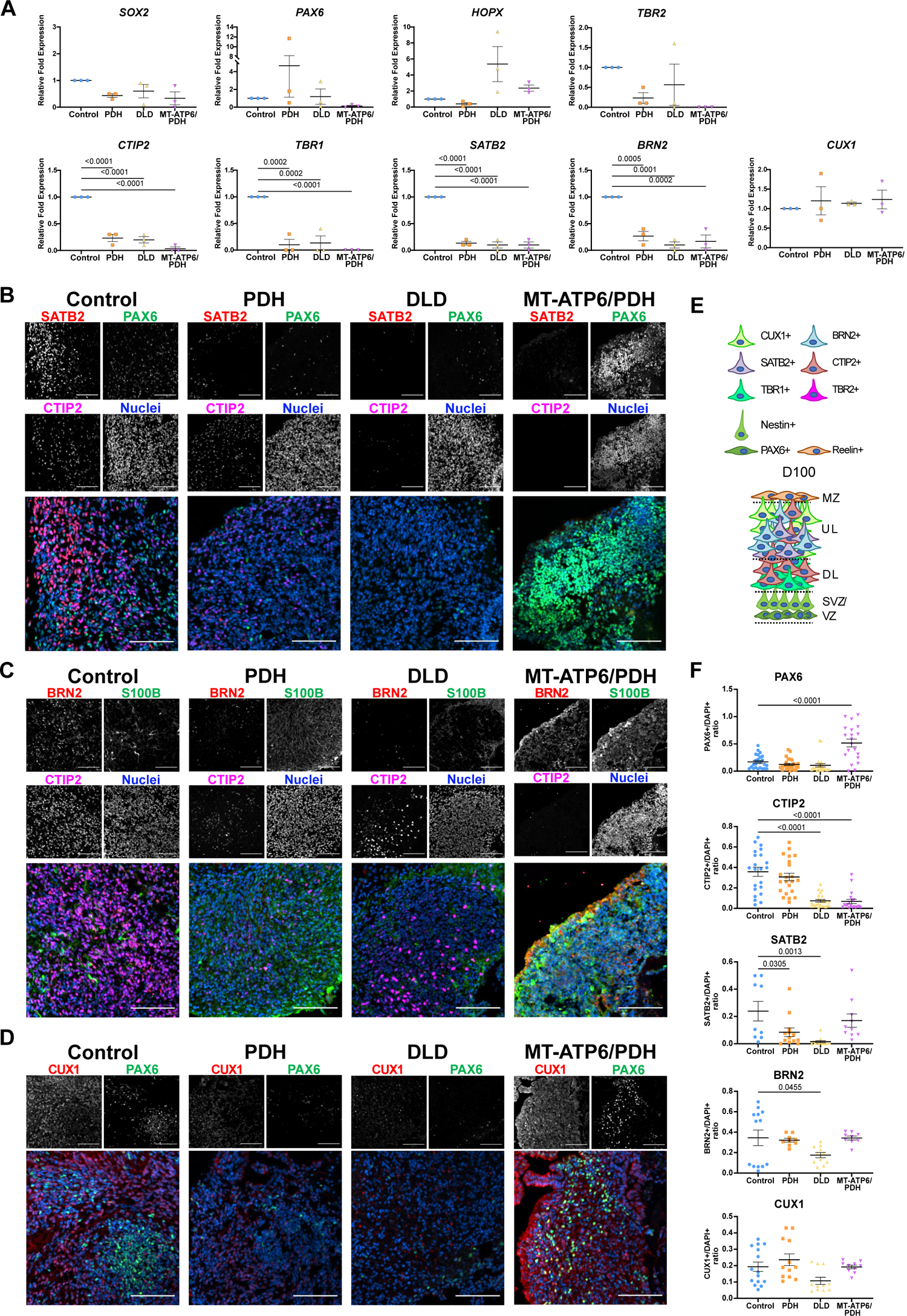
Leigh syndrome-derived brain organoids show defects in cortical layer formation at day 100. A. RT-qPCR quantification of the day 100 organoids. Neural progenitor cell populations were evaluated by the expression of *SOX2* and *PAX6*. Intermediate progenitor cells (IPCs) were identified with the marker *TBR2* and outer radial glia (oRG) was evaluated by the expression of *HOPX*. Markers *CTIP2*, *TBR1*, *SATB2*, *BRN2*, and *CUX1* were assessed for cortical development. B-D. Representative confocal images of day 100 brain organoids. LS-derived brain organoids present reduced expression of the upper layer markers SATB2 (B) and BRN2 (C) and deep layer marker CTIP2 (B & C). Expression of the astrocyte marker S100β was also observed in the cell lines (C). LS-derived brain organoids express the upper layer marker CUX1 and NPC marker PAX6 (D). Images were generated from at least three different organoids per genotype from independent organoid batches. Scale bar for B-D: 100μm. E. Schematic representation of the expected organization of the brain organoids at day 100. F. Quantification of day immunofluorescence staining for day 100 brain organoids. Upper layer marker SATB2 was significantly reduced in PDH mutant (p=0.0305). DLD mutant presented reduced expression of the cortical layer markers CTIP2 (p<0.0001), SATB2 (p=0.0013), and BRN2 (p=0.0455). The double mutant MT-ATP6/PDH shows a significant increase in the PAX6+ population (p<0.0001), as well as reduced expression of the cortical plate marker CTIP2 (p<0.0001). Data are shown as mean ±SEM. Comparisons were performed as ordinary one-way ANOVA with a Dunnett’s multiple comparisons test post-hoc. SVZ: subventricular zone, VZ: ventricular zone, DL: deep layers UL: upper layers, MZ: marginal zone.

To assess the proper cortical layer fate specification during development, we grew cerebral organoids until day 100 and probed for upper cortical layer markers (Figure 7) (Florio and Huttner, 2014; Lui et al., 2011; Saito et al., 2011). RT-qPCR analysis of the gene expression at this time point showed no significant differences in the expression of NPC markers *SOX2*, and *PAX6*; outer Radial Glia (oRG) marker *HOPX* and IPC marker *TBR2*. Major dysregulation was observed in the neuronal markers (Figure 5A). Cortical layer maker *CTIP2* (p<0.0001 in all cases), *TBR1* (p=0.0002 for PDH and DLD, p<0.0001 for MT-ATP6/PDH), *SATB2* (p<0.0001 in all cases) and *BRN2* (p=0.0005 for PDH, p=0.0001 for DLD and p=0.0002 for MT-ATP6/PDH) were significantly reduced in all LS genotypes at this timepoint. Pan-neuronal marker *TUBB3* (p=0.0013 for PDH, p=0.0002 for DLD, and p=0.0003 for MT-ATP6/PDH) was also reduced for all three mutants, suggesting a reduced capacity of commitment to a neuronal fate. Interestingly, the neuronal marker *CUX1* did not show significant differences in expression among cell types. While CUX1 is predominantly expressed in pyramidal neurons of the upper layers II-IV of the developing cortex (Leone et al., 2008; Nieto et al., 2004), its expression has been reported in the subventricular zone (Nieto et al., 2004) and cortical plate (Saito et al., 2011). It has also been reported as being co-expressed with PAX6+ and TBR2+ cells (Cipriani et al., 2015). Due to the reduced expression of the cortical and neuronal markers, the maintained expression of *CUX1* could reflect its conserved expression in NPC and IPC populations rather than in committed upper-layer neurons. Quantification of the immunofluorescence images (Figure 5B-F) showed that the late-born superficial layer marker SATB2 (layer IV) was reduced in PDH (p=0.0305) and DLD (p=0.0013) organoids (Figure 5B and 5F). Cortical layer III marker BRN2 was reduced in the DLD mutant (p=0.0455, Figure 5C and 5F). CTIP2+ cells were also reduced in DLD and MT-ATP6/PDH organoids (p<0.0001 in both cases. Figure 5B-C and 5F). On day 100, MT-ATP6/PDH organoids had a significant increase in PAX6+ cells (p<0.0001) compared to control (Figure 5B, 5C, and 5F). The reduction in the number and variety of cell types found in the LS organoids may explain the decrease in their diameter at day 100 (Supplemental Figure 4D).

As astrogliosis is a hallmark for LS (Lake et al., 2015), we looked at the expression of astrocyte markers at the mRNA and protein level. q-RT PCR analysis of neuronal marker *TUBB3* reveals a marked downregulation in all three lines (PDH p=0.0013, DLD p=0.0002, and MT-ATP6/PDH p=0.0003) that may be associated with the reduction in the number of cortical neurons. Astrocyte marker *SOX9* (Sun et al., 2017) did not show major differences in expression when compared to control. Analysis of other astrocyte markers such as Glial Fibrillary Acidic Protein *(GFAP*), S100 calcium-binding protein-β *(S100B*), Aldehyde Dehydrogenase Family 1 Member L1 (*ALDH1L1*), and *VIMENTIN* showed an increased, yet not significant, upregulation in some genotypes (Figure 6A). In the case of DLD, all the above-mentioned markers were increased compared to the control. The double mutant MT-ATP6/PDH had increased expression of *VIMENTIN* and *ALDH1L1*, while PDH showed an increment in *S100B* and *VIMENTIN*. At a protein level, DLD and PDH cerebral organoids showed increased staining of astrocyte markers GFAP and S100B, respectively, at day 100 (Figure 5C, 6B, and 6D). S100B was also increased in the double mutant, but not in a statistically significant manner. Staining for the pan-astrocyte marker Aldehyde Dehydrogenase-1 Family Member L1 (ALDH1L1) did not show major differences between the genotypes (Figure 6C and 6D). Immunofluorescence staining of the organoids for β3-TUBULIN showed a statistical difference between control and DLD (p=0.0174), but not in the other two mutants. The decrease in the diversity of neuronal cell types and the increase in the presence of S100β+ cells in the double mutants may suggest a switch to astrocyte fate during cortical development. Interestingly, the DLD organoids had higher GFAP staining when compared to control (p=0.0141) that may suggest an increase in the reactivity of the astrocyte population. Upregulation of the astrocyte markers GFAP, S100β, and VIMENTIN have been associated with astrocyte reactivity and in response to injury (Clarke et al., 2018; Escartin et al., 2021; Liddelow and Barres, 2017; Liddelow et al., 2017; Qi et al., 2017; Zamanian et al., 2012). Although the gene expression of these pan-reactive markers was not significant when compared to control, it may suggest activation of the glial population in response to the metabolic dysregulation in LS organoids.

**Figure 6.**
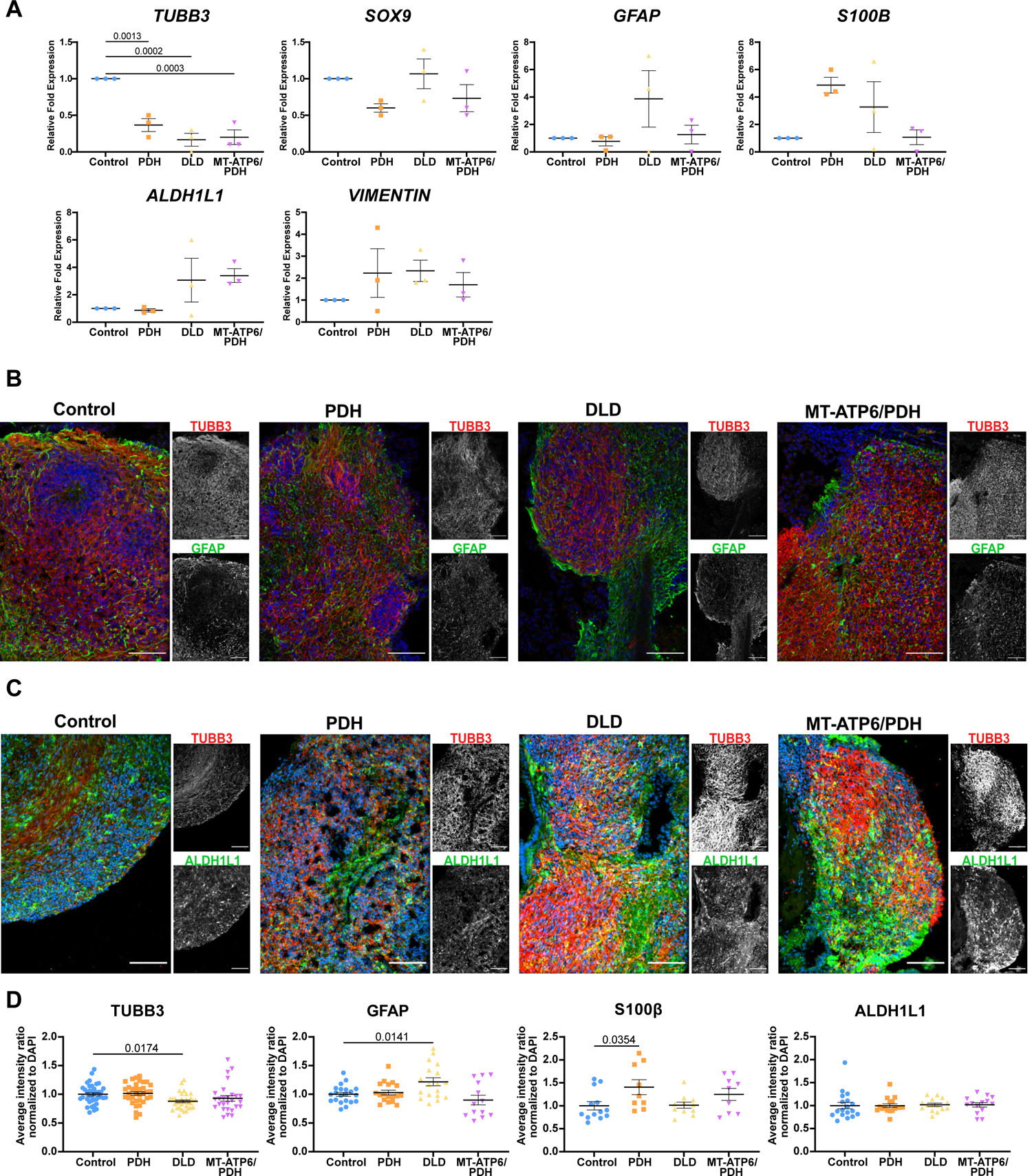
LS brain organoids show dysregulation of neuronal and glial markers at day 100. A. RT-qPCR analysis of neuronal and astrocytic genes at day 100. Neuronal marker *TUBB3* and astrocytic markers *SOX9*, *GFAP*, *S100B*, *VIMENTIN*, and *ALDH1L1* were evaluated. A significant decrease in the expression of the neuronal marker *TUBB3* was observed in all phenotypes. Fold change normalized to GPI and GAPDH as housekeeping genes. B-C. Day 100 immunofluorescence images of astrocytic markers GFAP (B) and ALDH1L1 (C), and neuronal marker βIII TUBULIN (TUBB3). Nuclei in the merged image correspond to the blue channel. Scale bar: 100μm. D. Immunofluorescence quantification of the neuronal and astrocytic staining. DLD mutant presented decrease staining in the neuronal marker TUBB3 (p=0.0174) and an increase in the astrocytic marker GFAP (p=0.0141); The PDH mutant shows a significant increase in the S100β+ population (p=0.0354). Scale bar for B-C: 100μm. Data are shown as mean ±SEM. Comparisons were performed as ordinary one-way ANOVA with a Dunnett’s multiple comparisons test post-hoc.

### Leigh syndrome-associated mutations disrupt the mitochondrial network in the ventricular zone of cerebral organoids

Mitochondria in murine NPCs form an elongated network (Khacho et al., 2016) which fragments as cells undergo neurogenesis (Iwata et al., 2020). Mitochondrial morphology was also evaluated in the VZ NPCs of the cerebral organoids. Cells positive for NPC marker SOX2+ demonstrated elongated mitochondrial networks that extend radially from the ventricle-like lumen (Supplemental Figure 6A). These results are significant because it is the first evidence demonstrating that the mitochondrial networks are remodeled in the developing human brain as reported in the developing mouse cortex (Iwata et al., 2020; Khacho et al., 2016). PDH mutant organoid NPCs have an increased axis length of their mitochondrial network in comparison with control (p=0.0078, Supplemental Figure 6B). As mentioned earlier, the stereotypical arrangement of the VZ was compromised in most MT-ATP6/PDH organoids. In the few areas where ventricle-like structures were identified with a conserved SOX2+ VZ, the mitochondrial network appeared more aggregated. This morphology was also observed in the clusters of SOX2+ cells that were scattered throughout the organoid. Quantification of the mitochondrial network for this mutant (Supplemental Figure 6B) showed differences in the mitochondrial network’s volume, diameter, surface area, and axis length (p<0.0001, in all cases). A higher volume and diameter may suggest an increase in mitochondrial aggregation in the VZ. Moreover, the difference in mitochondrial length may also correlate with the increased expression of *TBR2* observed by RT-qPCR (Iwata et al., 2020).

As NPCs generated by a dual SMAD monolayer method did not show major differences among the different genotypes, we looked at their mitochondria morphology to evaluate if the differences observed in the organoids were recapitulated in this setup. Characterization and quantification of various parameters of the mitochondrial network using structured illumination microscopy (SIM) showed that while control human NPCs showed elongation of the mitochondrial network, the DLD mutant shows an increase in mitochondrial number and decreased sphericity. Both DLD and PDH mutants have a significant increase in the number of branches in the network. (Supplemental Figure 6C-6D) which may reflect an increase in fusion events (Sukhorukov et al., 2012; Westrate et al., 2014). These parameters suggest that both the DLD and PDH mutant lines present alterations in mitochondrial morphology, which could be linked to the underlying changes in metabolic capacity (Rafelski, 2013) and developmental defects (Westrate et al., 2014).

### Metabolic dysregulation in Leigh syndrome-derived cerebral organoids

To explore early changes in metabolites, we perform metabolomic profiling of Day 40 organoids from control and LS samples. Metabolomic analysis showed 43 different metabolites that were statistically dysregulated in the LS organoids when compared to control (Supplemental Table 2 and Supplemental Figure 7). Out of these metabolites, 8 were dysregulated in PDH, 16 in DLD, and 32 in MT-ATP6/PDH (Figure 7A and Supplemental Figure 7).

**Figure 7.**
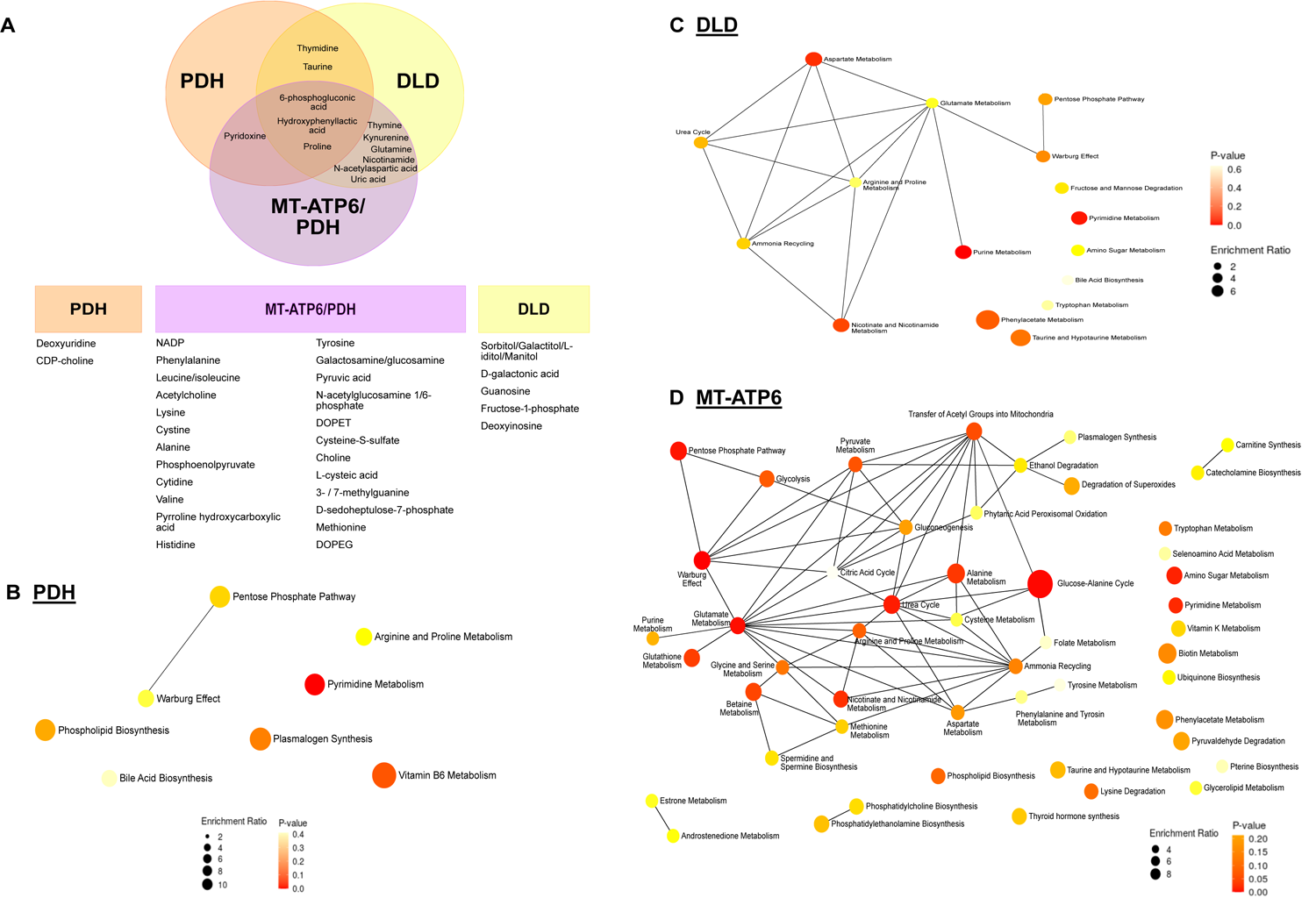
Day 40 LS organoids show changes in their metabolic profiles. A. 43 metabolites were statistically dysregulated (p<0.05 and FDR of 0.01) segregated by affected cell line. A total of three batches of 40-day organoids per line (4 independent organoids per line per batch) were analyzed as described in methods. B. Metabolite Set Enrichment Analysis for dysregulated metabolites enriched in the PDH mutant. C. Metabolite Set Enrichment Analysis for dysregulated metabolites enriched in the DLD mutant. D. Metabolite Set Enrichment Analysis for dysregulated metabolites enriched in the MT-ATP6/PDH mutant. Diameter of the node is determined by the level of enrichment, while the color of the node represents the p-value of the interaction.

The metabolites proline, 6-phosphogluconic acid, and hydroxyphenyllactic acid were dysregulated in all three LS mutants. Proline concentration was elevated when compared to control. High levels of proline have been associated with negative effects in brain function by interference in glutamatergic neurotransmission (Gogos et al., 1999; Vorstman et al., 2009). On the other hand, 6-phosphogluconic acid was reduced in all three LS cell lines. High concentrations of this metabolite have been associated with an active pentose phosphate pathway in early brain development in rats (Hakim et al., 1980) and its supplementation increased the diameter of neurospheres derived from the embryonic Ts1Cje mouse model of Down syndrome (Seth et al., 2020). In addition, hydroxyphenyllactic acid was elevated in DLD and PDH mutant organoids but downregulated in MT-ATP6/PDH. High levels of this metabolite have been reported in association with high lactate and pyruvate in pediatric lactic acidosis in patients with PDHc deficiency (Kumps et al., 2002; Stern, 1994).

Pyruvate was also increased in the MT-ATP6/PDH mutant organoids, correlating with the lactic acidosis expected in the organoids based on the patient phenotypes and the presence of the PDH mutation that hinders flux from pyruvate into the TCA cycle through acetyl-CoA. Moreover, the glycolysis/gluconeogenesis intermediate phosphoenolpyruvate was also elevated in these mutants. Increased levels of phosphoenolpyruvate in rat brains after ischemic injury are thought to have a protective role in cerebral ischemia *in vivo* (Geng et al., 2021) and in oxygen/glucose deprivation *in vitro* (Jiang et al., 2019).

Besides the previously mentioned metabolites, MT-ATP6/PDH mutant organoids presented increased levels of choline, cytidine, and leucine. Choline is a crucial metabolite for normal CNS development. Neural tube defects have been associated with a lack of choline during early pregnancy (Zeisel, 2006). It has also been shown to increase cell proliferation and decrease apoptosis in fetal rat hippocampal progenitor cells (Albright et al., 1999b, 1999a; Zeisel and Niculescu, 2006). Choline is also crucial for the production of the neurotransmitter acetylcholine, the sphingolipid sphingomyelin, and myelin (Oshida et al., 2003). Concomitantly, cytidine is used with choline for the generation of cytidine-5-diphosphocholine, a crucial intermediate in the biosynthesis of the cell membrane phospholipids phosphatidylcholine and phosphatidylethanolamine (Cansev, 2006; Rema et al., 2008). Increased abundance of the branched-chain amino acid leucine has been associated with the metabolic illness maple syrup urine disease and can be extremely neurotoxic (Bridi et al., 2005; García-Cazorla et al., 2014). This amino acid is considered ketogenic as its end products can enter the TCA cycle for energy generation or act as precursors for lipogenesis and ketone body production (Manoli and Venditti, 2016).

Pathway analysis of the dysregulated metabolites (Supplemental Tables 3, 4, and 5) show overlap in the pyrimidine metabolism, taurine, and hypotaurine metabolism, pentose phosphate pathway, arginine and proline metabolism, and aminoacyl-tRNA biosynthesis. Pyrimidine nucleotides are essential precursors for nucleic acid synthesis and are involved in polysaccharide and phospholipid biosynthesis, detoxification processes, and protein and lipid glycosylation (Fumagalli et al., 2017). Taurine and hypotaurine are osmotic regulators in the brain, as well as agonists to GABAergic and glycinergic neurons (Albrecht and Schousboe, 2005). Its presence in the developing brain is necessary for the correct development of axons and the formation of synaptic connections (Sturman, 1993). Dysregulation of the aminoacyl-tRNA biosynthesis pathways is well documented as causal etiology for several neurodevelopmental disorders such as leukoencephalopathies, microcephaly, and Leigh syndrome (Francklyn and Mullen, 2019; Ognjenović and Simonović, 2018).

We also performed metabolite set enrichment analysis (Figure 7B-C). Besides the previously mentioned pathways, the Warburg effect or aerobic glycolysis was also shared among the LS mutants. Although this effect is considered one of the hallmarks of cancer, it has also been associated with several homeostatic processes, including cell turnover and proliferation, and brain development (Bubici and Papa, 2019). Energy generation through aerobic glycolysis as a compensatory mechanism to overcome the metabolic deficiency in LS could suggest a survival adaptation of the cerebral organoids. Moreover, the shutoff of aerobic glycolysis is critical to neuronal differentiation in human NPCs. Inability to transition to neuronal oxidative phosphorylation causes apoptosis due to excessive conversion of pyruvate to lactate, and potentially a cell fate shift into GFAP-positive glial cells (Zheng et al., 2016b). Considering our observations that there is a marked deficit in MT-ATP6/PDH mutants to commit and generate neuronal subtypes and an increased signal in astroglial markers, these mutations may be impairing the ability to transition from aerobic glycolysis to OXPHOS as previously described with SURF mutations (Inak et al., 2019, 2021). The preferential switch to a glial fate may be promoted by astrocytes having low expression levels or lower active levels of the PDHα subunit (Bélanger et al., 2011; Halim et al., 2010; Itoh et al., 2003; Laughton et al., 2007).

Considering all the previous findings, the cerebral organoid system emerges as a promising model for the study of the effects of LS-associated mutations in brain development and the effects of metabolic impairment during corticogenesis.

## Discussion

Inborn errors of metabolism are rare genetic disorders resulting from defects in metabolic pathways (Agana et al., 2018; Das et al., 2010). Mitochondrial diseases are the most common group of inherited metabolic disorders and are among the most common forms of inherited neurological disorders (Gorman et al., 2016). These illnesses are challenging not only at the time of diagnosis but also during their medical evolution, as they involve multiple organ systems and have limited therapeutic options (Grier et al., 2018; Parikh et al., 2017; Schaefer et al., 2019).

Leigh syndrome is one of these rare inherited neurometabolic diseases, with more than 75 causal genes identified in both nuclear and mitochondrial DNA. It has an early onset, affecting most patients within their first year of life, although cases during teenage years and adulthood have been reported (Finsterer, 2008; Lake et al., 2016). As it is a highly heterogeneous disease, the establishment of animal and *in vitro* models has been challenging and limited to only select mutations. The few animal models available have been utilized for the development of therapeutic approaches with mixed results. Gene editing using adeno-associated virus in *Ndufs4*-/- mice has shown partial rescue of the phenotype (Di Meo et al., 2017). Supplementation of nicotinamide riboside to *Sco2*-/- mice showed improvement of the respiratory chain defect and increased exercise tolerance due to improved mitochondrial biogenesis (Cerutti et al., 2014). Hypoxia and low oxygen availability in the brain have also been shown to increase the life span and improve neurological findings in *Ndufs4*-/- mice (Ferrari et al., 2017; Jain et al., 2016, 2019).

Molecular testing and prenatal diagnosis of respiratory chain disorders using skin fibroblasts and muscle biopsies for diagnostic and research purposes are becoming mainstream procedures (Baertling et al., 2014; Calvo et al., 2006, 2012; Schubert and Vilarinho, 2020). Nevertheless, heteroplasmy in mitochondrial DNA among tissues can cause less pronounced or absent phenotypes in cultured cells (Baertling et al., 2014). Here we report the characterization and the subsequent generation of brain organoids from three commercially available Leigh syndrome fibroblast cell lines and age-matched control.

Three-dimensional differentiation generates higher numbers of NPCs and more mature neurons than two-dimensional differentiation (Chandrasekaran et al., 2017; Di Lullo and Kriegstein, 2017; Muratore et al., 2014; Paşca et al., 2015) in part due to an improved spatial cellular environment that influences cell fate specification. Tissue architecture, mechanical cues, cell-to-cell communication (Pampaloni et al., 2007), nutrient accessibility, oxygen tension, as well as morphogen gradients characteristic of 3D systems (Tibbitt and Anseth, 2012) aid to recapitulate the development of the central nervous system (CNS) up to approximately 16 weeks post-fertilization (Camp et al., 2015; Paşca et al., 2015). We observed that all the LS cerebral organoids failed to thrive at different time points. Although organoid development initially appeared normal in cell lines with nuclear-encoded LS mutations, at later time points the overall diameter decreased, presumably due to failure to generate upper-layer neurons. This reduction of late-born neurons may be due to the overall reduction in the PAX6+ NPCs in both PDH and DLD compared to controls. Interestingly, PDH cerebral organoids displayed arrested growth very early on, correlating with the microcephaly observed in the proband.

Clinical data from LS patients include marked gliosis as part of the characteristic findings (Baertling et al., 2014, 2016; Lake et al., 2015; Schubert and Vilarinho, 2020). While this marked gliosis is potentially associated with a reactive process secondary to neuronal damage, an intriguing alternate possibility is that progenitor cells may have an increased propensity to differentiate down the astrocyte lineage due to LS-causative mutations and mitochondrial-associated dysregulations. Previous studies have shown that reactive astrocytes acquire molecular hallmarks of radial glial cells. It was also shown through genetic fate mapping that mature astroglial cells can dedifferentiate and resume proliferation (Robel et al., 2009, 2011). Thus, the increase in the glial-specific marker S100β in PDH organoids and DLD multipotency cultures, as well as in GFAP staining in DLD organoids and the upregulation trend in the mRNA expression of the astrocyte markers could either reflect that chronic metabolic stress induced by Leigh syndrome mutations activates a brain injury response, or that the inhibition of mitochondrial metabolism in neural progenitor cells could cause defects in lineage selection (Escartin et al., 2021). Due to the differential expression and activation levels of the PDH complex in astrocytes (Bélanger et al., 2011; Halim et al., 2010), a predisposition of these cell lines to commit to an astroglial fate cannot be ruled out. Also, longer culture (>100days) of these organoids may allow for the interrogation of a gliosis phenotype in more detail. Analysis of A1 specific reactivity markers may point out if the upregulation of *GFAP, S100B, ALDH1L1,* and *VIMENTIN* is associated with a neuroinflammation type of response that has been associated with neuronal damage (Escartin et al., 2021; Liddelow et al., 2017).

The formation of lesions in LS has been described as the result of OXPHOS dysfunction and subsequent ATP depletion. Neuronal dysfunction is suspected to trigger chronic gliosis (Baertling et al., 2016). In patients, the gliosis phenotype can be accompanied by vascular hypertrophy and the production of excess ROS, which increases neuronal damage (Lake et al., 2015). However, due to the lack of vascularization in the organoid model, replicating the vascular abnormalities associated with LS is not feasible in this system.

In a previous study (Hattori et al., 2016), the metabolic signature analysis of iPSCs derived from a mitochondrial encoded LS mutation (m.10191T>C) showed differences in the abundance of pyruvate and lactate, among others. Interestingly, the metabolic difference between control and LS cells was reported to revert to normal after EB differentiation (Hattori et al., 2016). In our study, metabolomic analysis from organoids shows that the observed changes in the metabolites are in line with the clinical observations of LS patients. Changes in blood and cerebral spinal fluid concentration of lactate and pyruvate are common diagnostic tools for LS (Hattori et al., 2016) and other mitochondrial diseases (Barshop, 2004; Buzkova et al., 2018; Esterhuizen et al., 2017; Rahman and Rahman, 2018). While changes in the NADH/NAD+ ratio, *de novo* nucleotide synthesis, and in other metabolites from the ETC complex III and TCA cycle, were also identified, these were modest considering the genetic alterations in the mutant cell lines should directly affect these pathways. This could point out to metabolic compensatory mechanisms that may be engaged during development. Moreover, the disruption in the metabolic network observed in the LS cerebral organoids correlates with the severity and mortality of the disease in the probands. Although aerobic glycolysis was identified as a significantly affected pathway in all the mutants, the effects of the MT-ATP6/PDH mutation reflected the importance of competent glycolysis to OXPHOS transition in the early brain development (Zheng et al., 2016b).

The metabolic dysregulation of the affected tissues in LS may have a direct effect on mitochondrial morphology and function. Mitochondrial fragmentation is a hallmark of glycolytic cell types such as stem cells and cancer cells (Chen and Chan, 2017; Rastogi et al., 2019). Moreover, neurogenesis defects have been observed in the context of mitochondrial morphology dysregulation and are considered to be upstream regulators of self-renewal and cell fate decisions in stem cells (Iwata et al., 2020; Khacho et al., 2016). In addition, energetic requirements have been shown to directly impact the capacity of progenitor cells to migrate and thrive in 3D environments (Zanotelli et al., 2018, 2019). Hence, mitochondrial morphology disruption observed in the double mutant MT-ATP6/PDH organoids is congruent with the metabolic and developmental profiles of these mutant organoids. To our knowledge, this is the first time that mitochondrial morphology in the cortex has been analyzed in a human model system of LS brain development and highlights the critical function of mitochondrial network plasticity for the proper specification of cell fate and survival. Studies in murine brains demonstrated a period of plasticity in postmitotic cells where mitochondrial morphology determines the fate of the daughter cells (Iwata et al., 2020). Our results also highlight the direct link of mitochondrial morphology with early events in human brain development.

The profound dysregulation of corticogenesis in the double mutant MT-ATP6/PDH may suggest that some pregnancies harboring LS-causing mutations may not be viable, which could lead to an underestimation of the prevalence of the disease in the population (Feeney et al., 2019). Prenatal genetic evaluation is now being performed for cases where there is a known risk for mitochondrial mutations (Craven et al., 2017; White et al., 1999). This testing can be performed as early as 10-12 weeks post conception and is usually requested if there is a history of a previously affected child or first-degree relative (Nesbitt et al., 2014). Interpretation of these tests is challenging in the case of mitochondrial mutations due to heteroplasmy; the biopsied tissues may exhibit a different mutational burden compared with other fetal tissues (Ferlin et al., 1997; Harding et al., 1992; Nesbitt et al., 2014; Steffann et al., 2007).

Taken together, our study sheds new light on the morphological and functional LS alterations impacting early events of neurogenesis. We were able to identify new genetic alterations in LS samples by using whole-exome sequencing and mitochondrial DNA sequencing. We described the effects of LS mutations on early development, underscoring the critical function of metabolism in human neurogesis. Our works also provides a comprehensive phenotypic characterization of available patient samples to encourage their utilization as model systems for uncovering the mechanisms underlying neuronal cell death in the context of LS and as human platforms for drug discovery.

## Detailed Methods

### Key Resource Table

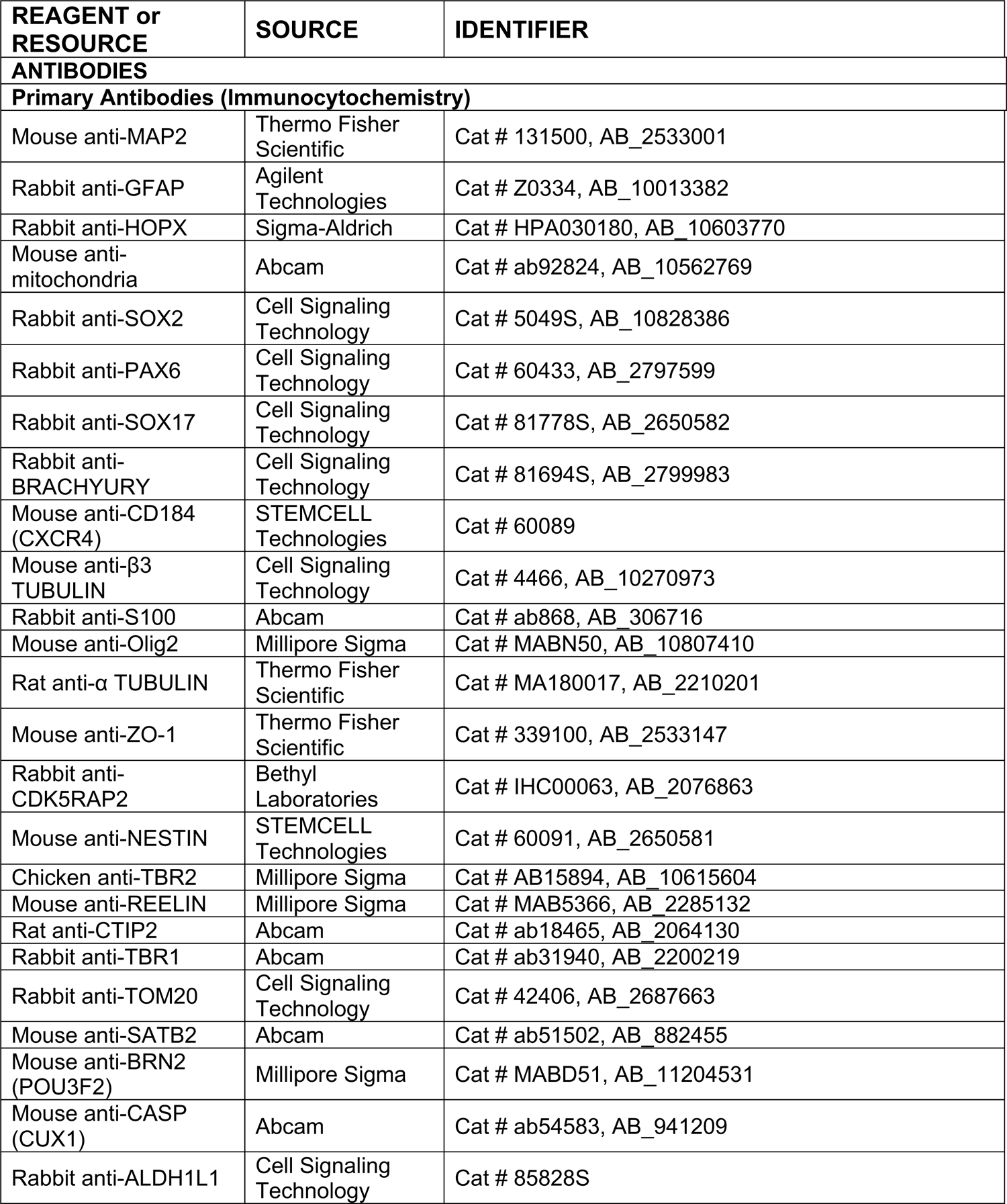
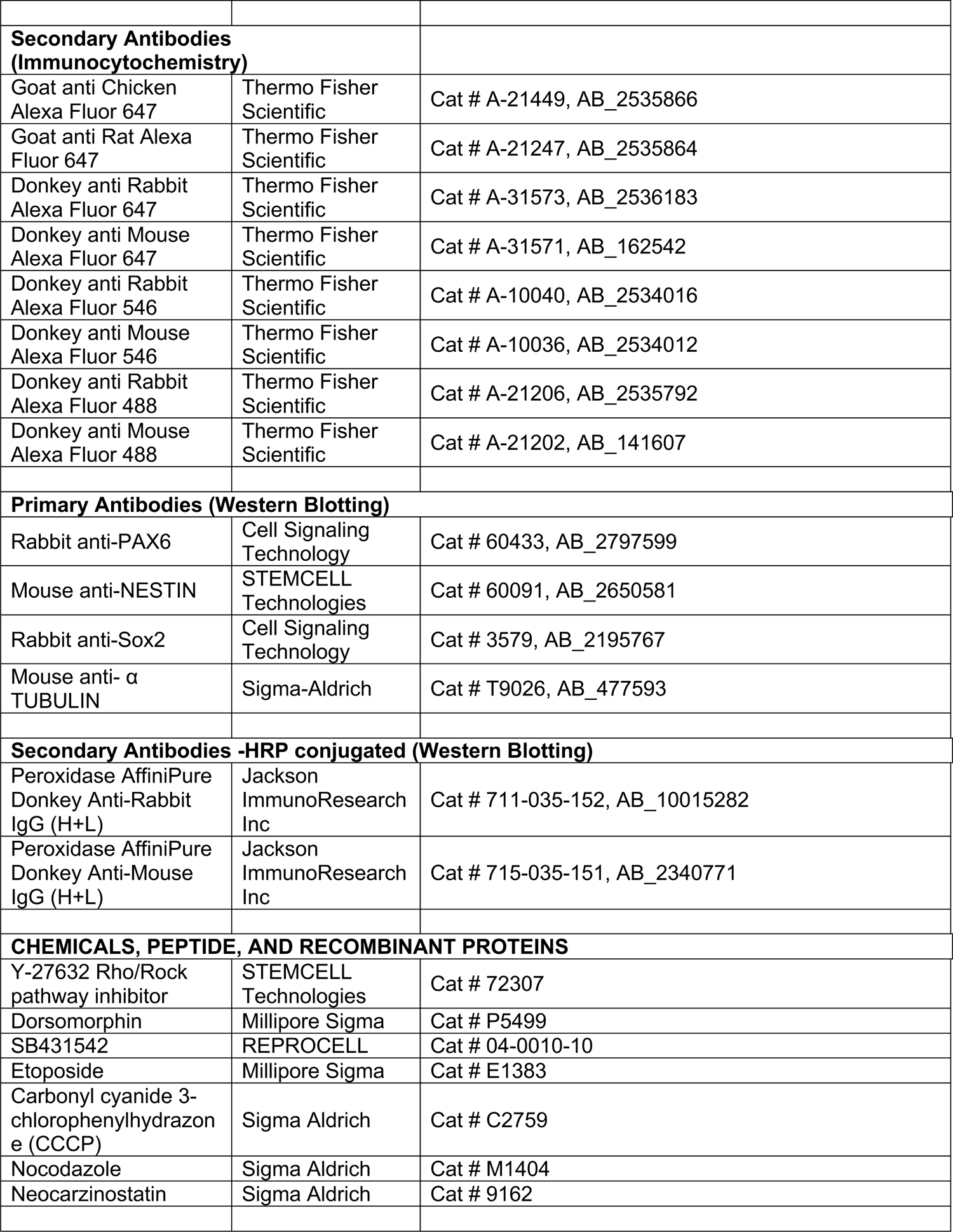
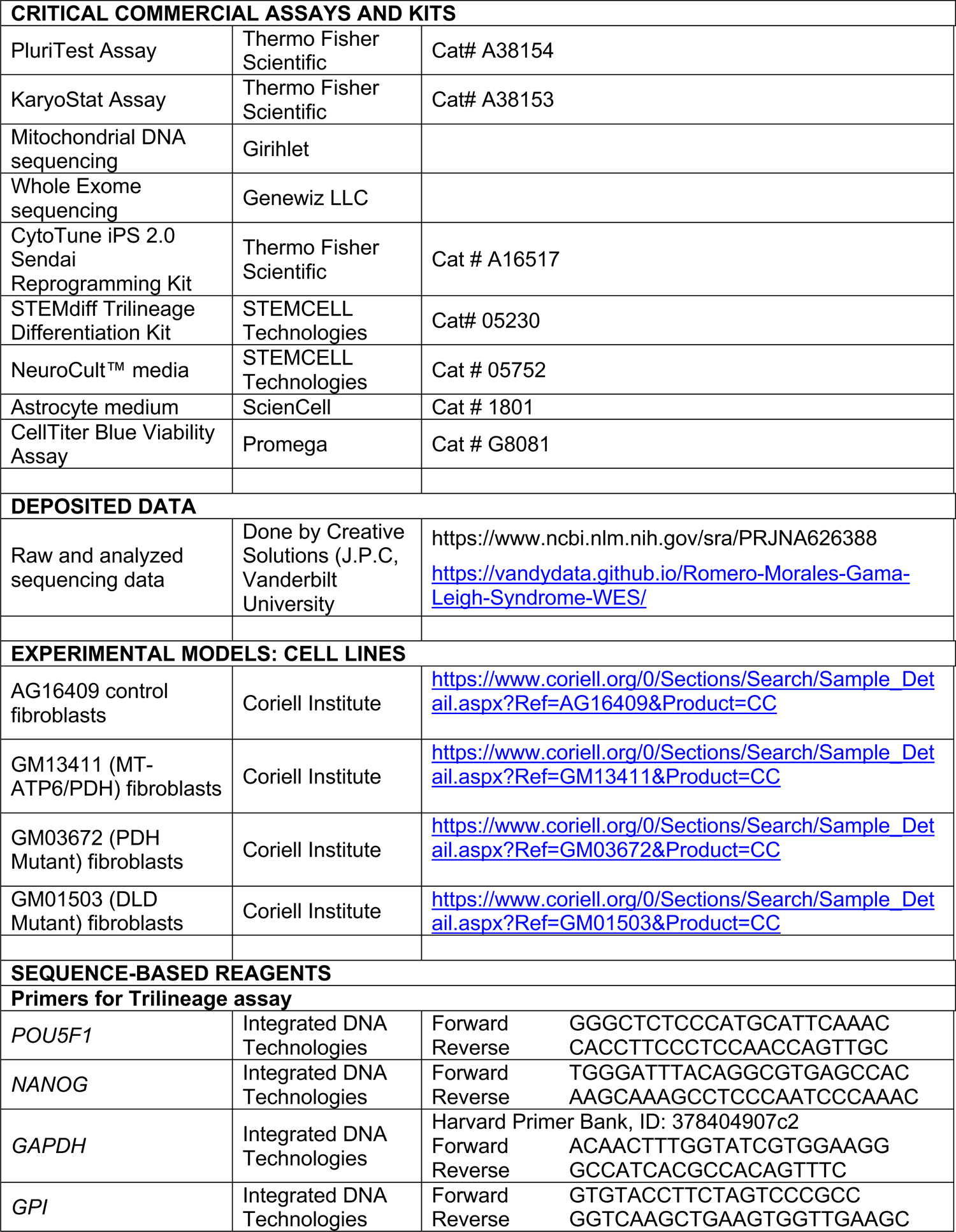
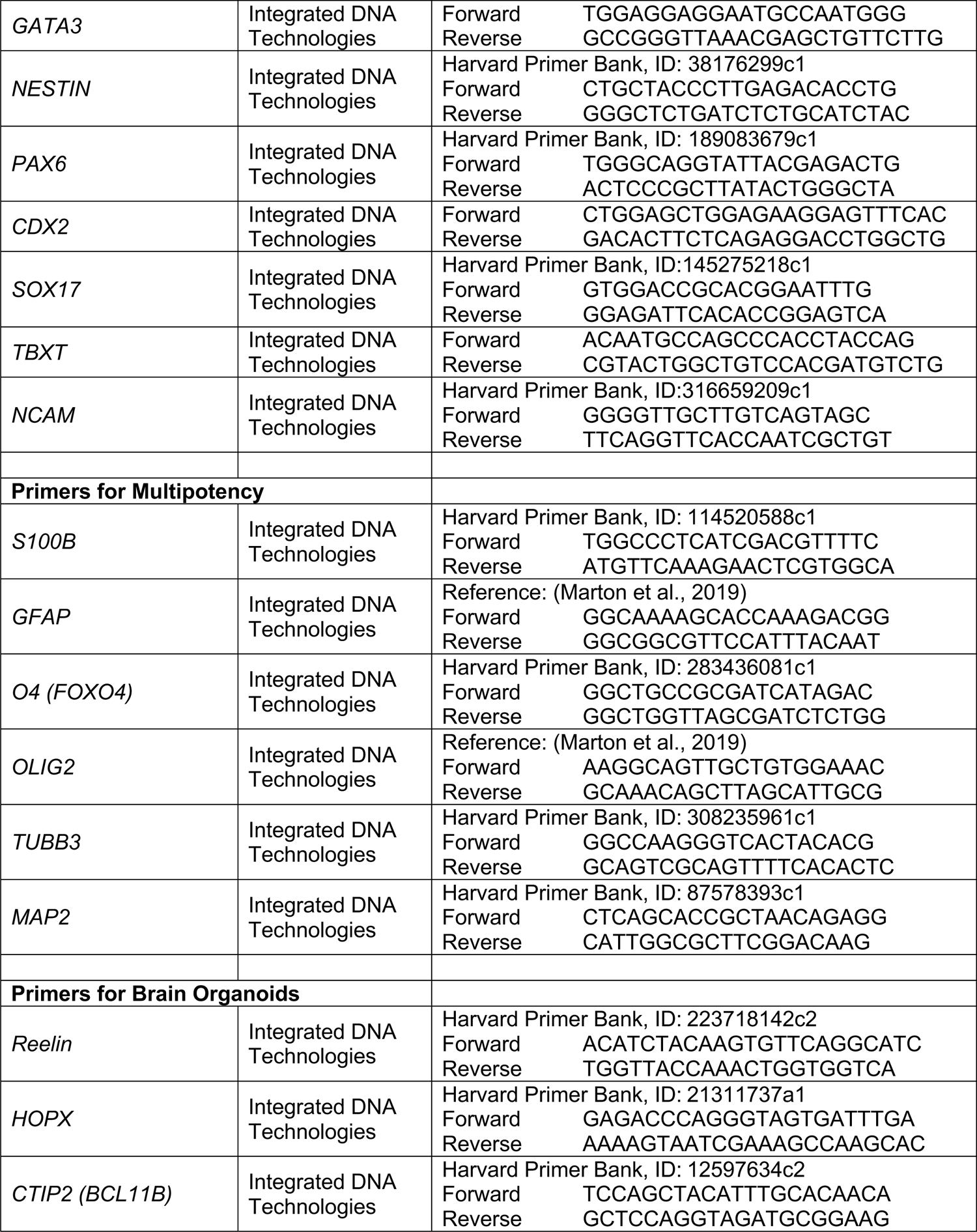
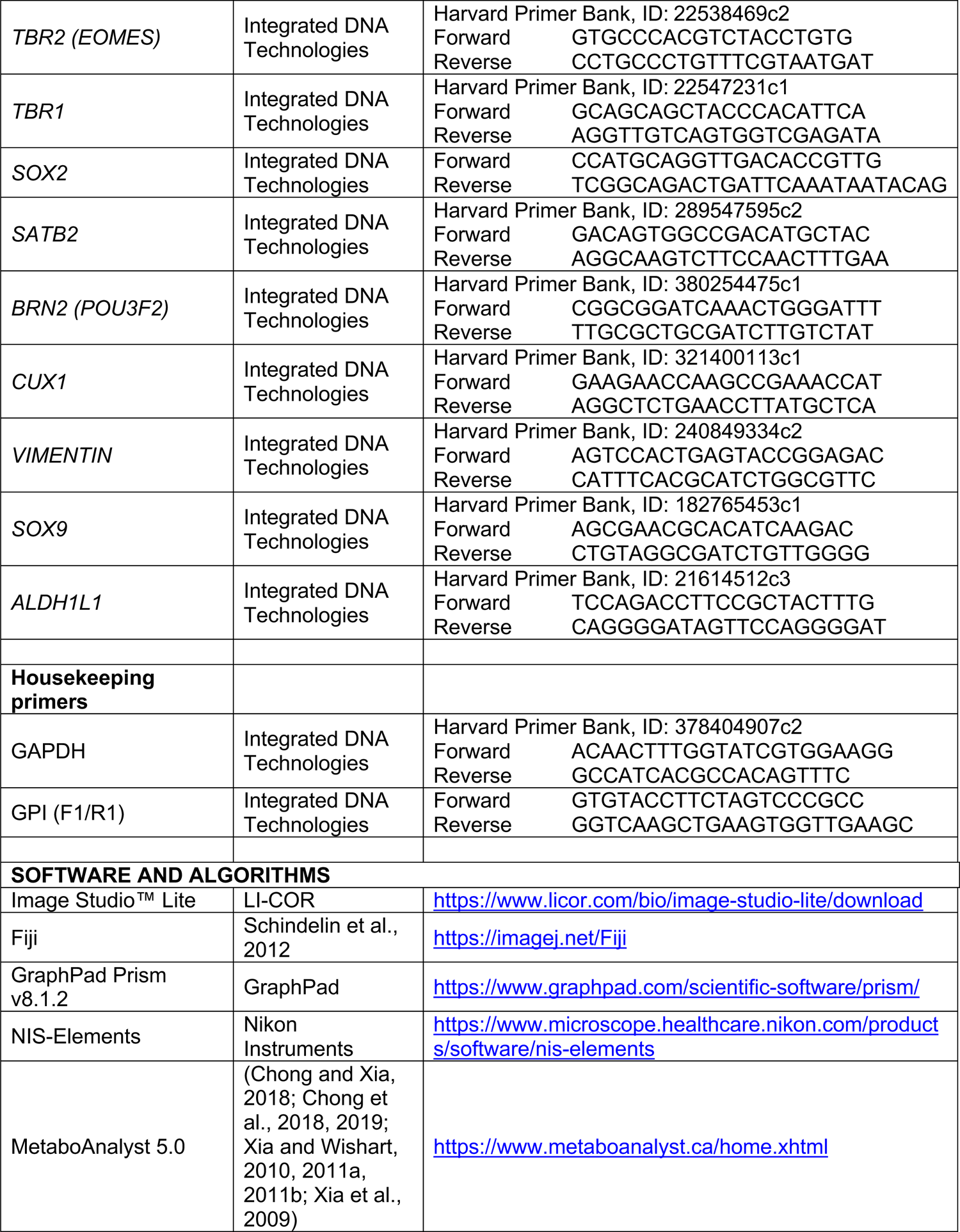
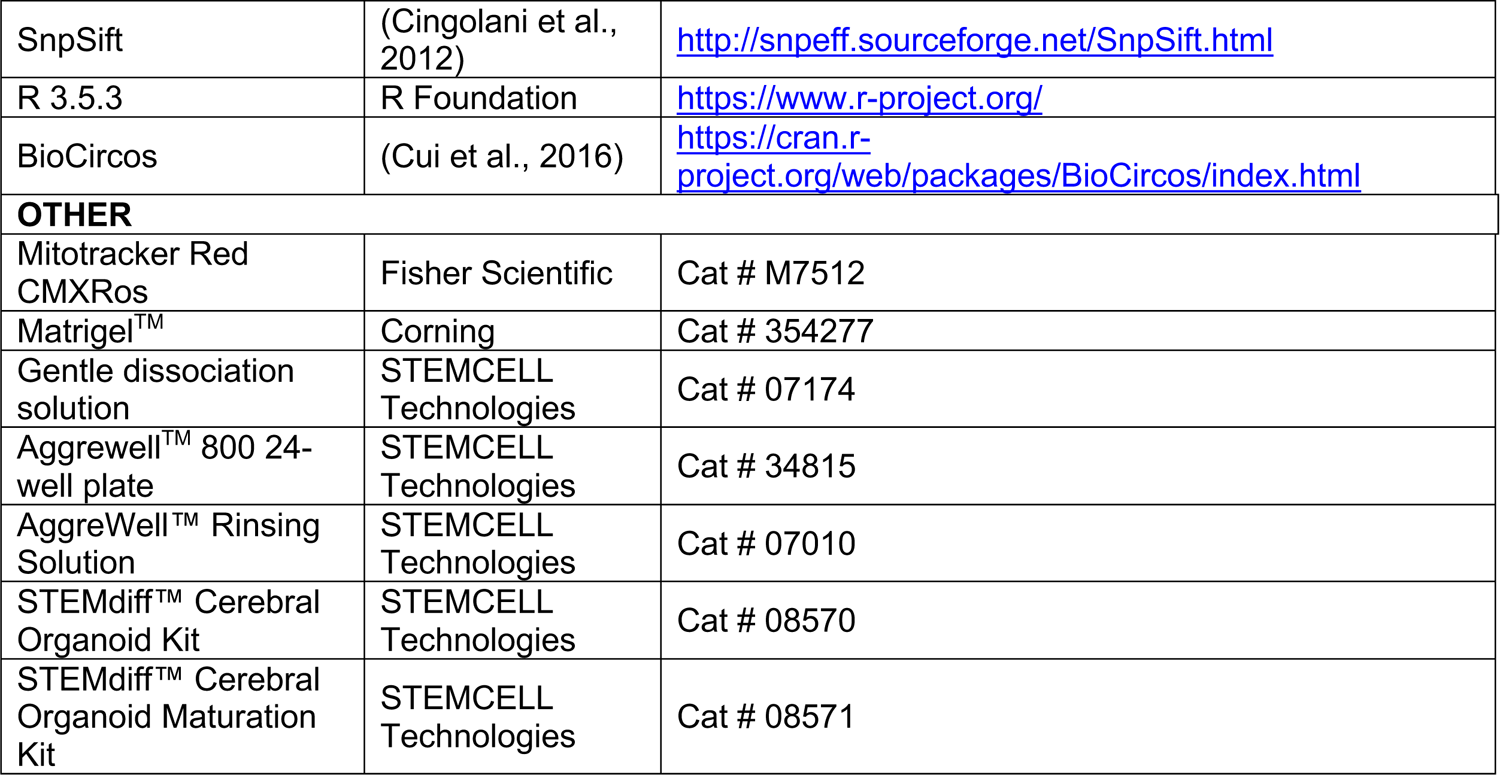

### Lead Contact and Materials Availability

Further information and requests for resources and reagents should be directed to the Lead Contact, Vivian Gama.

### Experimental Model and Subject Details

The Coriell cell line IDs were as follows: GM01503, GM03672, GM1341. Information about the Leigh syndrome cell lines used in this study can be found in Supplementary Table1. Control skin fibroblast cell line AG16409 was also obtained from Coriell Institute, Philadelphia, PA. The donor was a 12-year-old apparently healthy Caucasian male. Cells were negative for Mycoplasma. Fibroblasts were maintained in Dulbecco’s Modified Eagle Medium: Nutrient Mixture F-12 DMEM/F-12 (Gibco cat # 11330032) supplemented with 10% Fetal bovine serum (Sigma cat# F2442) in 100mm cell culture plates (Eppendorf, cat # 0030702115) in a 37°C 5% CO2 incubator.

## Method Details

### Whole Exome sequencing

Fibroblast cell pellets from each cell line (>1 million cells) were shipped on dry ice for whole-genome exome sequencing to Genewiz, Plainfield, NJ. The Illumina HiSeq-X was used to perform 150nt paired-end sequencing.

### Mitochondrial sequencing

Fibroblast cell pellets from each patient (>1 million cells) were shipped on dry ice for mitochondrial sequencing to Girihlet Inc. Oakland, CA. The sequencing configuration used was 80bp single-end sequencing, 20 million reads per sample.

### hiPSC Generation and Characterization

Human fibroblasts were purchased from healthy control and patients (Coriell Institute, Philadelphia, PA, USA). Induced pluripotent stem cells were derived from human fibroblasts using Sendai virus-based reprogramming kit (CytoTune-iPS Sendai Reprogramming Kit; cat #. A13780-01; Thermo Fisher), according to manufacturer’s instructions. After 3-4 weeks, 2-3 colonies per sample were transferred to fresh 6-well plates and were expanded and gardened for 3 passages before freezing. All iPSC cell lines were maintained in E8 medium in plates coated with Matrigel (Corning, cat # 354277) at 37°C with 5% CO2. Culture medium was changed daily. Cells were checked daily for differentiation and were passaged every 3-4 days using Gentle dissociation solution (STEMCELL Technologies, cat # 07174). All experiments were performed under the supervision of the Vanderbilt Institutional Human Pluripotent Cell Research Oversight (VIHPCRO) Committee.

### Analysis of Pluripotency

The pluripotency of each iPSC clone was determined using a microarray-based tool known as PluriTest (Thermo Fisher; cat# A38154) as an alternative to the teratoma assay. Samples were outsourced to Thermo Fisher for PluriTest and further analysis. Low passage iPSC cell pellets (>1 million cells) were frozen and shipped on dry ice. Additionally, the expression of pluripotency genes *POU5F1* and *NANOG* was assessed by qPCR.

### Analysis of Chromosomal abnormalities

The presence of any chromosomal abnormalities in the newly generated iPSCs was determined using a microarray-based tool known as KaryoStat (Thermo Fisher; cat# A38153) as an alternative to chromosomal G-banding. Samples were outsourced to Thermo Fisher for KaryoStat and further analysis. Low passage iPSC cell pellets (>1 million cells) were frozen and shipped on dry ice to Thermo Fisher.

### Trilineage differentiation

The STEMdiff Trilineage differentiation kit (STEMCELL Technologies, cat# 05230) was used to functionally validate the ability of newly established iPSCs to differentiate into three germ layers, as per the manufacturer’s instructions. Single-cell suspensions of 2×10^6^ cells/well, 5×10^5^ cells/well, 2×10^6^ cells/well were seeded for ectoderm, mesoderm, and endoderm, respectively, in their corresponding medium at day 0 in 6 well plates. The cultures were maintained for 7 days, 5 days, and 5 days for ectoderm, mesoderm, and endoderm, respectively. The differentiation was assessed by immunocytochemistry and qPCR.

### NPC differentiation and multipotency characterization

For monolayer differentiation of the iPSCs into NPCs, cells were dissociated into single cells using Gentle Cell Dissociation Reagent (STEMCELL Technologies, cat # 07174) for 8 minutes at 37°C. Live cell counts were performed using Trypan blue (0.4%) staining (Invitrogen, cat # T10282) using a Countess™ Automated Cell Counter. Cells were then seeded in a Matrigel-coated 6-well plate (Eppendorf, cat # 0030720113) to 2.5×10^6^ cells/well with dual SMAD inhibitor media supplemented with Dorsomorphin (1μM) and SB431542 (10μM) (Chambers et al., 2009) supplemented with ROCK inhibitor. Daily media changes were performed and passaging of the cells was done every 7-9 days. Cells for NPC marker analysis were collected at the end of the first 9 days of differentiation.

Mitochondrial imaging was performed by fixing the NPCs at 90% confluency and staining with anti-mitochondria (Abcam, cat #ab92824, 1:200 dilution). Structured Illumination Microscopy (SIM) was accomplished in 3D-SIM mode on a Nikon Instruments N-SIM, equipped with an Apo TIRF 100x SR 1.49NA objective, DU-897 EMCCD camera (Andor), 405nm and 561nm lasers. Images presented herein are maximum intensity projections after image stacks were first acquired (5 phase shifts and 3 rotations of diffraction grating, 120nm/axial step via piezo) and subsequent stack reconstruction in NIS-Elements software (Nikon Instruments, Inc.). Other than linear intensity scaling, no further image processing was performed post-reconstruction.

For multipotency analysis, culture media was changed to NeuroCult™ media and maintained for 4 weeks. Samples were then fixed and stained for neuron and oligodendrocyte markers. Astrocyte differentiation was performed by seeding on a Matrigel-coated plate 1.5×10^6^ cells/cm^2^ (TCW et al., 2017). The following day, the media was changed to Astrocyte medium (ScienCell, cat # 1801) and maintained for 20 days. Full media changes were done every 2 days. Samples were then fixed and stained for an astrocyte marker. Images were acquired with a Nikon Instruments Ti2 inverted fluorescence widefield microscope equipped with a Plan Apo Lambda 20X 0.75 NA objective, DS-Qi2 camera (Nikon Instruments), and X-Cite 120LED light source (Excelitas). The differentiation was also assessed by qPCR.

### Neural rosette differentiation

To generate neural rosettes, we dissociated the cells into a single-cell suspension and seeded 3.0×10^6^ cells/well of an Aggrewell^TM^ 800 in dual SMAD inhibitor media. EBs were incubated at 37°C with 5% CO2, with minimal disruption during the first 48 hours. Media changes, 50-75% of the total volume, were performed every 2 days. On Day 5, EBs were harvested according to the manufacturer protocol and transferred to a 35mm imaging plate (Cellvis, cat # D35-14-1.5-N) coated with Matrigel. Daily media changes were performed up to day 9 when cells were fixed with 100% ice-cold Methanol (Fisher Scientific, cat # A454-4). Images were acquired on a Nikon Instruments Ti2 inverted fluorescence microscope, equipped with a Yokogawa X1 spinning disk head, Andor DU-897 EMCCD, Plan Apo Lambda 0.75 NA 20X objective for representative figures, and Plan Fluor 0.45 NA 10X objective for neural rosette quantification, piezo Z-stage, as well as 405-,488-,561-, and 647 -nm lasers. Acquisition and analysis were performed using NIS-Elements. Neural rosette quantification was accomplished by scripting a segmentation-based image analysis routine to detect, enumerate, and measure rosette lumen area based on the ZO-1 signal. Briefly, max intensity projections of each field were generated, followed by GPU-based denoising of the resulting image. Intensity-based thresholding was then applied based on criteria established for ZO-1 signal segmentation using control images. Restrictions on resultant binaries were implemented to throw out binaries intersecting image borders, morphometries deviating severely from rosette associated geometries, as well as for those not meeting minimum size requirements. This routine could be run in batch across many image stacks to increase the sample size and robust nature of the data. Measured data was exported to Excel for further analysis.

### Cerebral Organoids

Cerebral organoids were generated as described in (Romero-Morales et al., 2019) with some modifications. Briefly, organoids were generated using the STEMdiff™ Cerebral Organoid Kit (STEMCELL Technologies; Cat# 08571, 08570). iPSCs were dissociated into single cells using Gentle Cell Dissociation Reagent (STEMCELL Technologies, cat # 07174) for 8 minutes at 37°C. Homogeneous and reproducible EBs were generated by using a 24-well plate AggreWell™ 800 (STEMCELL Technologies, cat # 34815). On Day 7, high-quality EBs were embedded in Matrigel (Corning, cat # 354277). On Day 10, the Matrigel coat was broken by vigorously pipetting up and down and the healthy organoids were transferred to a 60mm low attachment culture plate (Eppendorf, cat # 003070119). The plates were then moved to a 37°C incubator and to a Celltron benchtop shaker for CO2 incubators (Infors USA, cat # I69222) set at 85rpm. Full media changes were performed every 3–4 days. Transmitted-light images were acquired using an EVOS® XL Core Imaging System. The software used for processing was ImageJ. c. For qPCR, all organoids were pooled together for RNA extraction. Day 40 cerebral organoids were utilized for metabolomics, 4 organoids per genotype were run and analyzed individually.

### Organoid tissue preparation and Immunohistochemistry

Tissue preparation was performed as described in (Romero-Morales et al., 2019). Briefly, organoids were fixed in 4% Paraformaldehyde in Phosphate Buffered Saline (PBS), washed 3 times with PBS, and then incubated in 30% sucrose solution overnight at 4°C. Organoids were embedded in 7.5% gelatin/10% sucrose solution (Sigma, catalog G1890-100G and S7903-250G) and sectioned with a cryostat (Leica CM1950) at 15um thickness. For immunostaining, slides were washed with PBS before permeabilization with 0.2% Triton-X in PBS for 1 hr. Tissues were blocked with blocking medium consisting of 10% donkey serum in PBS with 0.1% Tween-20 (PBST) for 30 min. Incubation with primary and secondary antibodies was done using standard methods. Confocal images of the organoids were acquired using the aforementioned spinning disk microscope with Plan Fluor 10X 0.45 NA and Plan Apo Lambda 0.75 NA 20X objectives (macrostructures) and Apo TIRF 1.49 NA 100X objective (mitochondria imaging). NIS-Elements software was used for image acquisition and rendering.

### RNA Extraction and Synthesis of cDNA

Cells cultured in 6 well plate, were collected after a wash with PBS, using 600μl Trizol reagent. The samples were spun down at 12,000 g after the addition of 130μl of chloroform and incubated at room temperature for 3 minutes. The aqueous phase of the sample was collected 200μl at a time until reaching the edge of phase separation. RNA precipitation was done by incubating with 300μl of isopropanol for 25 minutes, followed by centrifugation at 12,000 g for 10 min at 4°C. The RNA pellet was washed with ethanol, semi-dried, and resuspended in 30μl of DEPC water. After quantification and adjusting the volume of all the samples to 1μg/μl, the samples were treated with DNAse (New England Biolabs, cat # M0303). 10μl of this volume was used to generate cDNA using the manufacturer’s protocol (Thermofisher, cat#4368814).

For RNA isolation from brain organoids, the same protocol mentioned above was followed with the volumes adjusted for 1mL of Trizol.

### Quantitative RT PCR (RT-qPCR)

1ug of cDNA sample was used to run RT-qPCR for the primers mentioned in the table. QuantStudio 3 Real-Time PCR machine, SYBR green master mix (Thermo Fisher, cat#4364346), and manufacturer instructions were used to set up the assay.

### Immunocytochemistry

Cells were fixed with 4% paraformaldehyde (Electron Microscopy Sciences, cat # 15710-S) in PBS for 20 min at 4°C. Blocking and permeabilization were done in 5% donkey serum (Jackson ImmunoResearch Inc, cat # 017-000-121) + 0.3% Triton X-100 (Sigma Aldrich, cat # T9284) in TBS for 1 hr at room temperature. After this, cells were treated with primary and secondary antibodies using standard methods. Cells were mounted in Vectashield (Vector Laboratories, cat # H-1000) prior to imaging.

### Western Blotting

Cultured cells were lysed in 1% Triton buffer containing PMSF (ThermoFisher Scientific, cat # 36978), PhosSTOP (Roche, cat # 4906837001), and protease inhibitor cocktail (Roche, cat # 4693132001). Protein concentrations were determined using the bicinchoninic acid (BCA) method (Thermo Scientific, cat # 23227). Gel samples were prepared by mixing 30μg of protein with LDS sample buffer (Life Technologies, cat # NP0007) and 2-Mercaptoethanol (BioRad, cat # 1610710) and boiled at 95°C for 5 minutes. Samples were run on 4-20% Mini-PROTEAN TGX precast gels (BioRad, cat # 4561096) and transferred onto polyvinylidene difluoride (PVDF) membrane (BioRad, cat # 1620177) overnight at 4°C. Membranes were blocked in 5% milk in TBST prior to primary antibody incubation. Antibodies used for Western blotting are described in the Key Resource table.

### Cell titer blue assay

After the 24-h exposure to individual treatments of 50μM etoposide, 80μM CCCP, 100ng/mL nocodazole, and 5ng/mL neocarzinostatin, 20 μl of Cell Titer Blue reagent from Cell Titer Blue assay (Promega, cat # G8081) was added to each well of 96 well plate. Background fluorescence was calculated by adding 10% Triton in PBS to some wells. The fluorescence generated by the reduction of resazurin to resorufin by live cells was measured using a Beckman coulter DTX 880 multimode plate reader (Beckman Coulter, Brea, California) (570/600 nm).

### Metabolomics analysis

Day 40 brain organoids, at least 4 individual organoids per genotype, were collected rinsed with ice-cold sterile 0.9% NaCl and flash-freeze in liquid nitrogen. For metabolite extraction, cells were resuspended in 225uL of cold 80% HPLC grade methanol/20% HPLC grade water per 1×10^6^ cells. After resuspension, cells were flash-frozen in liquid nitrogen and thawed rapidly in a 37°C water bath 3 times. Next debris was removed by centrifugation at max speed in a tabletop microcentrifuge at 4°C for 15 min. Metabolite-containing supernatant was transferred to a new tube, dried, and resuspended in 50% acetonitrile while the pellet was used for protein quantification. Samples were analyzed by Ultra-High-Performance Liquid Chromatography and High-Resolution Mass Spectrometry and Tandem Mass Spectrometry (UHPLC-MS/MS).

Specifically, the system consisted of a Thermo Q-Exactive in line with an electrospray source and an Ultimate3000 (Thermo) series HPLC consisting of a binary pump, degasser, and auto-sampler outfitted with an Xbridge Amide column (Waters; dimensions of 4.6mm × 100mm and a 3.5μm particle size). Mobile phase A contained 95% (vol/vol) water, 5% (vol/vol) acetonitrile, 10mM ammonium hydroxide, 10mM ammonium acetate, pH = 9.0; and mobile phase B was 100% Acetonitrile. The gradient was as follows: 0 min, 15% A; 2.5 min, 30% A; 7 min, 43% A; 16 min, 62% A; 16.1-18 min, 75% A; 18-25 min, 15% A with a flow rate of 400μL/min. The capillary of the ESI source was set to 275°C, with sheath gas at 45 arbitrary units, auxiliary gas at 5 arbitrary units, and the spray voltage at 4.0kV. In positive/negative polarity switching mode, an m/z scan range from 70 to 850 was chosen, and MS1 data was collected at a resolution of 70,000. The automatic gain control (AGC) target was set at 1×10^6^ and the maximum injection time was 200 ms. The top 5 precursor ions were subsequently fragmented, in a data-dependent manner, using the higher energy collisional dissociation (HCD) cell set to 30% normalized collision energy in MS2 at a resolution power of 17,500. Data acquisition and analysis were carried out by Xcalibur 4.1 software and Tracefinder 4.1 software, respectively (both from Thermo Fisher Scientific). The peak area for each detected metabolite was normalized by the total ion current which was determined by the integration of all of the recorded peaks within the acquisition window.

Normalized data was uploaded to MetaboAnalyst (https://www.metaboanalyst.ca/home.xhtml) for analysis. Samples were normalized to control, and a one-way ANOVA was performed to compare between the groups. Fisher’s least significant difference method (Fisher’s LSD) was performed as a post-HOC comparison. Enrichment and pathway analysis was also performed using this platform.

### Bioinformatic Analysis

Bioinformatic analysis began with Variant Call Format (VCF) files provided by GENEWIZ (see *Whole Exome sequencing* section above), both for SNP and indels. SnpSift version 4.3t (Cingolani et al., 2012) was used to process and filter these files for downstream analysis. Details extracted included gene symbol, Entrez gene ID and name, UniProt ID, Ensembl ID, chromosome and position, reference variant, alternative variant, quality of the call, allele name, type of SNP, impact of the SNP, and the genotype of each sample. From these filtered outputs, we generated SNP/indel reports that allowed us to look at sample-specific SNPs and indels, as well as perform aggregate-level functions for grouping and statistical analysis.

To generate the SNP/indel circular chromosome plots, the top 20 genes that had variants in all three samples were plotted, ranked by frequency of variants per gene. The outside track is used to visualize the chromosomes and marked gene locations. For each sample, we used a single track to show the variant frequency as a circular scatter plot, with the height of the scatter points representative of the variant quality metric, which is a Phred-scaled probability that a REF/ALT polymorphism exists at the variant site. Similarly, for SNPs in the mitochondrial chromosome, we used the same approach for visualization.

### Quantification and Statistical Analysis

No statistical methods were used to pre-determine sample sizes. All experiments were performed with a minimum of 3 biological replicates unless specified. Statistical significance was determined by unpaired Student’s t-test or by one- or two-way ANOVA as appropriate for each experiment. GraphPad Prism v8.1.2 was used for all statistical analysis and data visualization.

Error bars in all bar graphs represent the standard error of the mean or standard deviation as described for each Figure, while scattered dot plots were represented with boxes (with median and SD) and whiskers (minimum and maximum values).

For neural rosette experiments, ROI were randomly selected using the nuclear (DAPI) staining channel. Images were processed with NIS Elements software with our Neural rosette lumen identification Macro. Outliers were removed from the neural rosette area analysis as post-processing quality control for the NIS Element macro using GraphPad Prism v8.1.2. ROUT (Robust regression and Outlier removal) method was used with a False Discovery Rate of 1%. For cerebral organoid experiments, 4 independent batches were generated. At time points day 30 and day 100, at least 5 organoids per cell line were collected. Immunofluorescence images of at least 3 independent organoids were acquired per condition slide. Image processing was done by NIS Elements and Fiji software. For the organoid quantification, images were processed with NIS Elements using the General Analysis 3 tool. 3D thresholding macros were generated for each set of slides and quantified by either bright spot count (nuclear staining) or mean intensity of the ROI. For the day 30 mitochondria analysis, the ROUT (Robust regression and Outlier removal) method was used with a False Discovery Rate of 1%.

Organoid efficiency evaluation was performed on day 10 using 4X transmitted-light images acquired using an EVOS® XL microscope. Two observers were blinded to the cell line identifier and counted the number of normal and defective (no epithelial buds or more than 75% of the area is not developed) organoids. Criteria for normal and defective organoids was based on (Lancaster and Knoblich, 2014).

### Software and Data Availability

All raw data in FastQ format for whole-exome sequencing and mitochondrial sequencing have been deposited to the Short Read Archive as BioProject PRJNA626388, available at https://www.ncbi.nlm.nih.gov/sra/PRJNA626388. All source code and documents are available via https://vandydata.github.io/Romero-Morales-Gama-Leigh-Syndrome-WES/.

## Author contributions

A.R.M. and V.G. designed experiments, interpret the data, and wrote the manuscript. A.R.M., G.L.R., A.R., and M.L.R. performed experiments and analyzed data with technical support of H.T. J.P.C. and L.H. analyzed the genomic data, generated the corresponding figures, and the associated method section. P.M.A provided pluripotency data analysis and technical support for data analysis. B.M generated the neural rosette analysis macro for NIS Elements. N.S.C., G.S.M., and R.P.C. provided technical support, led the experimental design and data analysis for metabolomics and its corresponding method section.

## Conflict of interests

The authors declare no competing interests.

## Acknowledgments

We thank Dr. Nicholas Mignemi (Vanderbilt Nikon Center for Excellence) for his technical support with image acquisition and processing. We would like to thank members of the Gama Laboratory for helpful discussions and comments on the manuscript. Funding was provided by the following grants: 1R35GM128915-01 (to VG); 1R21 CA227483-01A1 (to VG); the Precision Medicine and Mental Health Initiative sponsored by the Vanderbilt Brain Institute (to VG). Image acquisition and analysis were performed in part through the use of the Nikon Center of Excellence within the Vanderbilt Cell Imaging Shared Resource (supported by NIH grants CA68485, DK20593, DK58404, DK59637, and EY08126), Vanderbilt University Medical Center’s Translational Pathology Shared Resource supported by NCI/NIH Cancer Center Support Grant 2P30 CA068485-14, and the Vanderbilt Mouse Metabolic Phenotyping Center Grant 5U24DK059637-13. Whole exome sequencing and mitochondrial sequencing results were analyzed by Creative Solutions. Metabolite measurements were performed by the Northwestern University RHLCCC Metabolomics Core (Dr. Peng Gao) and were supported by the following NIH grants to N.S.C.: NIH2PO1HL071643-11A1, NIH1R35CA197532-01, NIH1PO1AG049665-01, NIH/NCI grant to G.S.M: T32CA09560, and Northwestern University Pulmonary and Critical Care Department’s Cugell Predoctoral Fellowship to R.P.C.

## Supplemental Information

**Supplemental Figure 1, Related to Figure 1.**
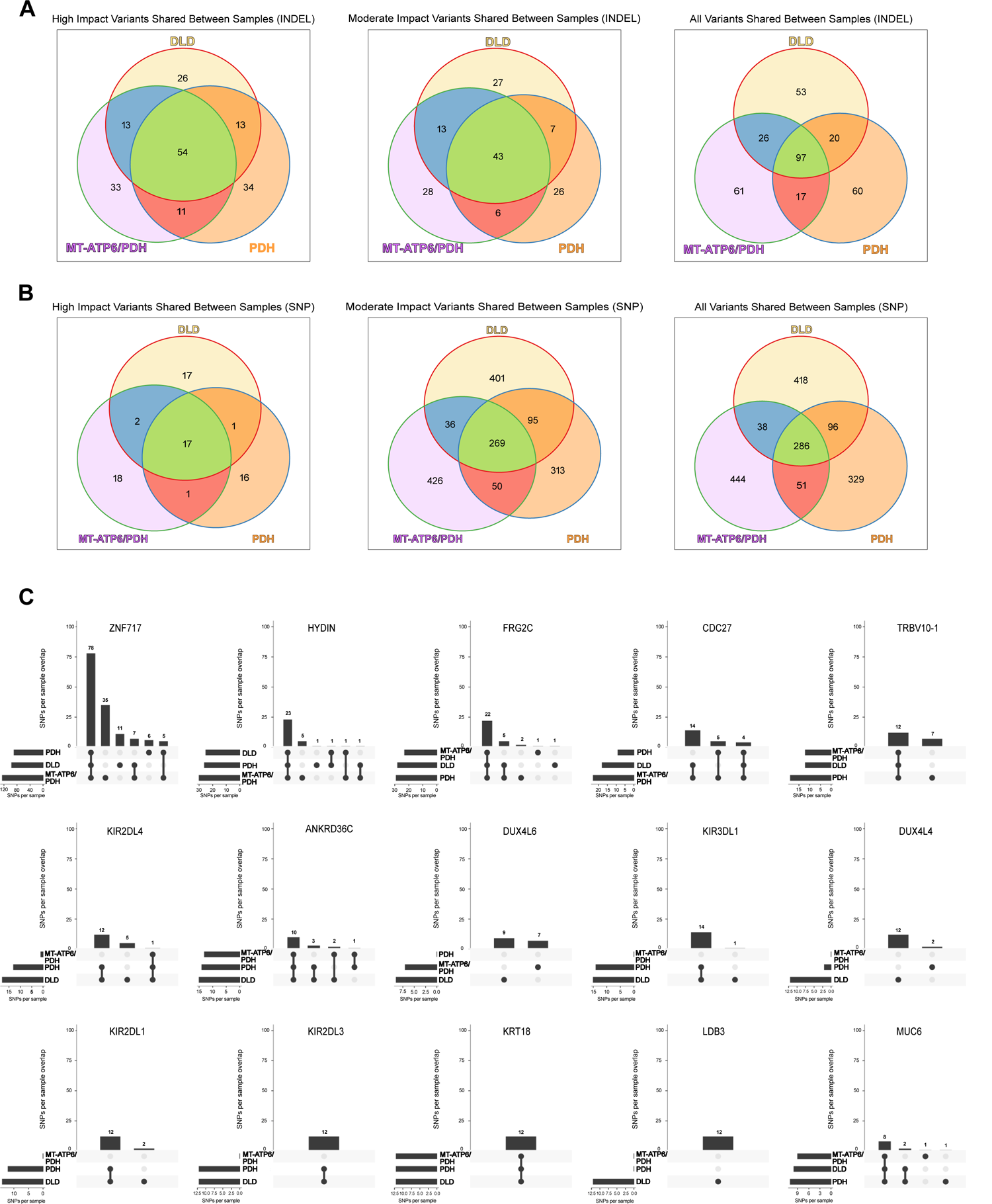
A-B. Venn diagram of the distribution of high, moderate and all impact annotations across samples for INDELs (A) and SNPs (B). Pie charts of the annotation distributions within samples. C. Top 15 genes with high impact SNP variants identified in the WES analysis. Number of overlapping SNPs per sample are denoted in as vertical bars, while the number of SNPs present in each phenotype are noted in the horizontal bars. *ZNF717*: Zinc finger protein 717, *HYDIN*: HYDIN axonemal central pair apparatus protein, *FRG2C*: FSHD region gene 2 family member C, *CDC27*: Cell division cycle 27, *TRBV10-1*: T cell receptor beta variable 10-1, *KIR2DL4*: Killer cell immunoglobulin like receptor, two Ig domains and long cytoplasmic tail 4, *ANKRD36C*: Ankyrin repeat domain 36C, *DUX4L6*: Double homeobox 4 like 6, *KIR3DL1*: Killer cell immunoglobulin like receptor, three Ig domains and long cytoplasmic tail 1, *DUX4L4*: Double homeobox 4 like 4, *KIR2DL1*: Killer cell immunoglobulin like receptor, two Ig domains and long cytoplasmic tail 1, *KIR2DL3*: Killer cell immunoglobulin like receptor, two Ig domains and long cytoplasmic tail 3, *KRT18*: Keratin 18, *LDB3*: LIM domain binding 3, *MUC6*: Mucin 6.

**Supplemental Figure 2, Related to Figure 2.**
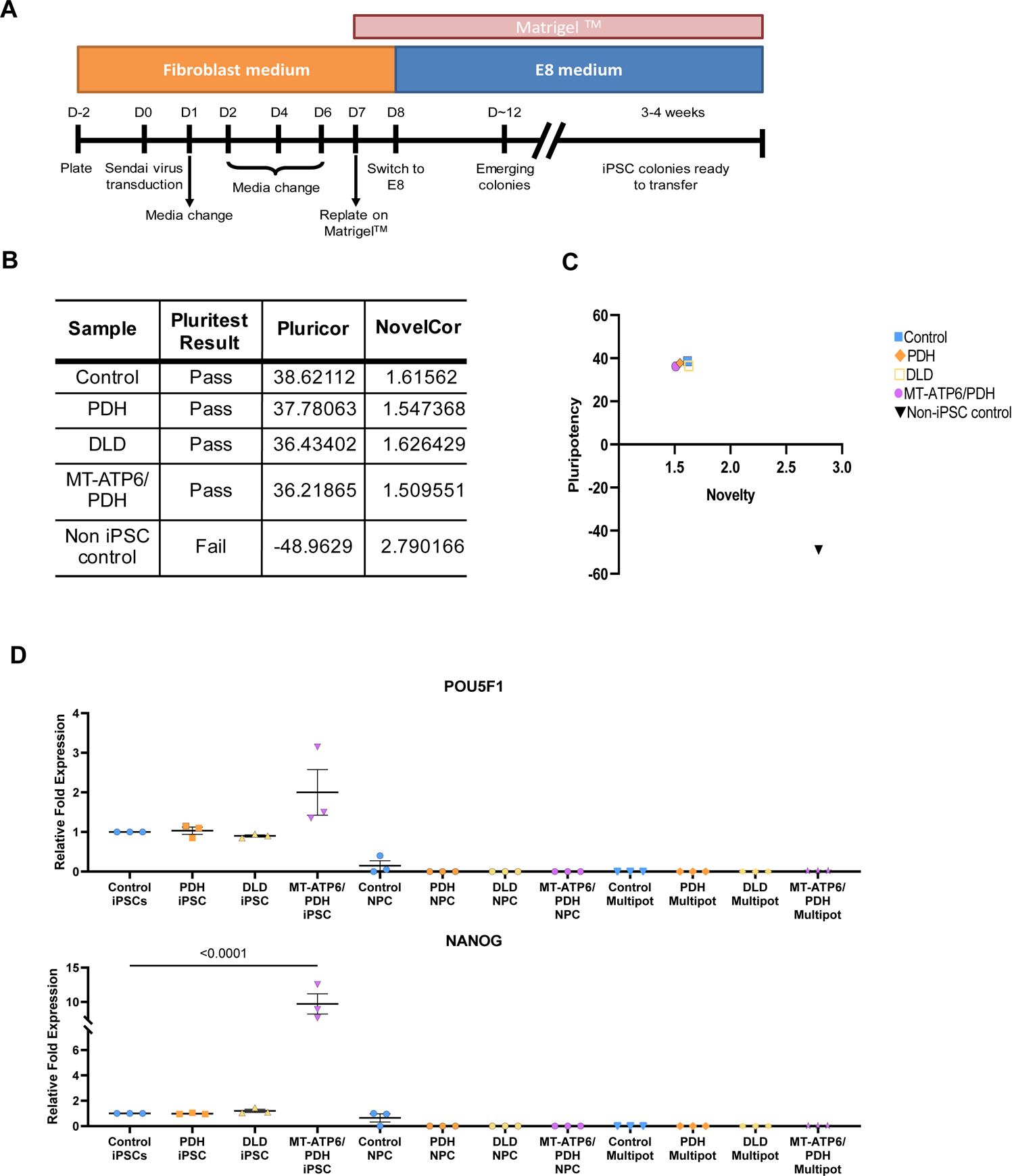
Characterization of Leigh syndrome iPSCs. A. Schematic representation of the fibroblast reprogramming protocol. B-C. Pluripotency characterization of the LS iPSCs. Samples were analyzed against samples in a reference data set (The International Stem Cell Initiative, 2018) (B). The distribution of the samples compared with a non-iPSC control shows clustering of the samples in the high pluripotency and low novelty quadrant (C). D. qPCR for the pluripotency genes *POU5F1* and *NANOG (*p < 0.001). Fold change normalized to GPI and GAPDH as house-keeping genes. iPSC: induced pluripotent stem cells, NPC: neural progenitor cells, Multipot: neural multipotency differentiation.

**Supplemental Figure 3, Related to Figure 3.**
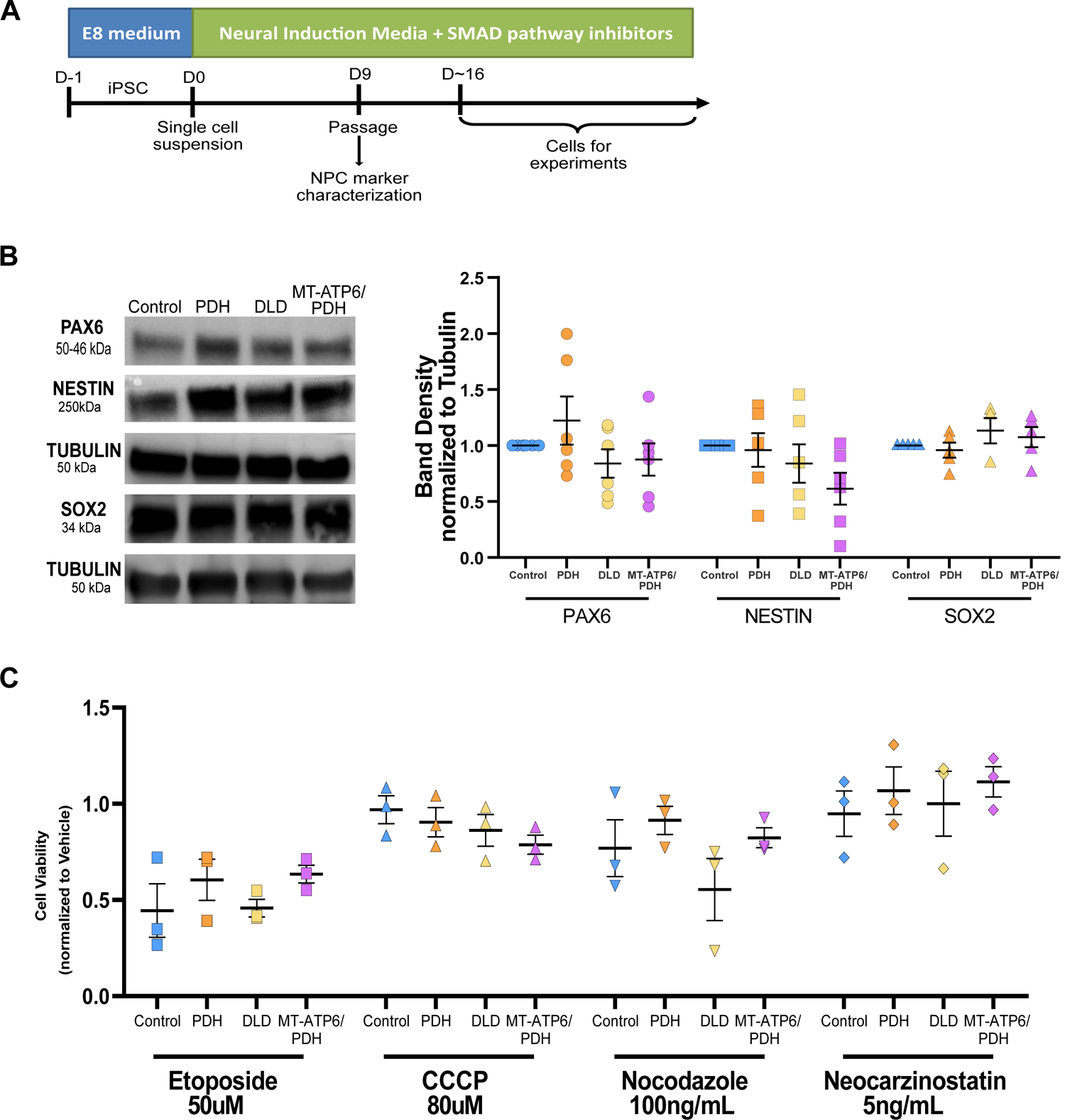
Leigh syndrome derived NPCs are multipotent and do not present increased sensitivity to pharmacological stressors. A. Schematic of two-dimensional neural differentiation. B. Immunoblot of protein expression of neural markers PAX6, NESTIN, and SOX2 (Left). Quantification of protein expression of neural markers PAX6, NESTIN and SOX2. Band density normalized to TUBULIN as a loading control (Right). C. Leigh syndrome NPCs do not show enhanced sensitivity towards different stressors. LS NPCs have similar cell viability compared to control when exposed to the DNA damaging agents etoposide and neocarzinostatin, the mitochondrial toxicant CCCP, and the microtubule destabilizer nocodazole.

**Supplemental Figure 4.**
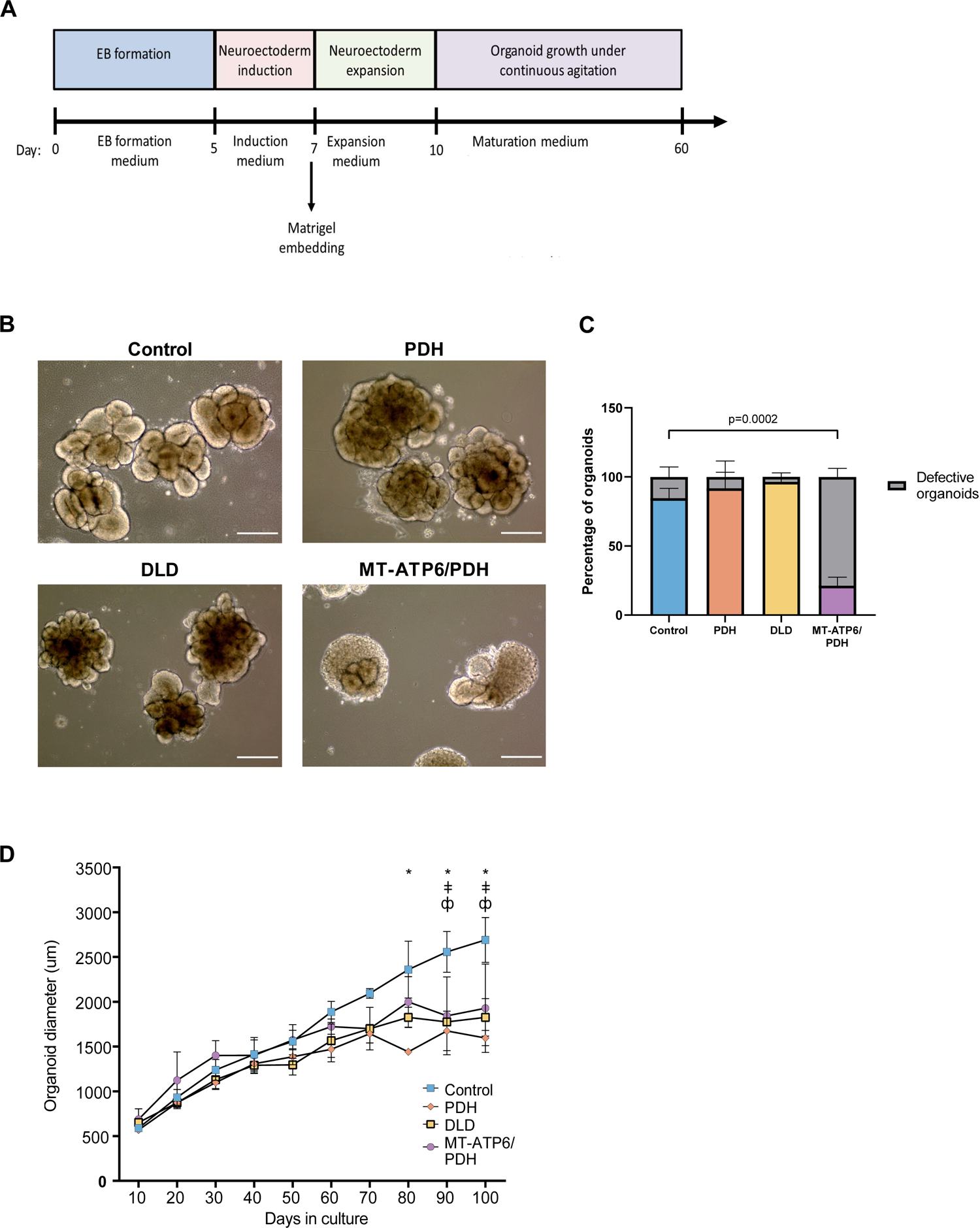
MT-ATP6/PDH brain organoids display defective differentiation at day 10. A. Schematic of the brain organoid generation protocol. B. Brightfield images (4X) of day 10 brain organoids. MT-ATP6/PDH shows disorganized cellular growth that do not resemble neuroepithelial buds. Scale bar: 300μm. C. Quantification of the defective organoids at day 10 by cell line. D. Brain organoid growth curves. LS derived brain organoids show reduction in size when grown for 100 days. * p < 0.05 for Control and PDH mutant; ǂ p < 0.05 for Control and DLD mutant, ф p < 0.05 for Control and MT-ATP6/PDH mutant.

**Supplemental Figure 5.**
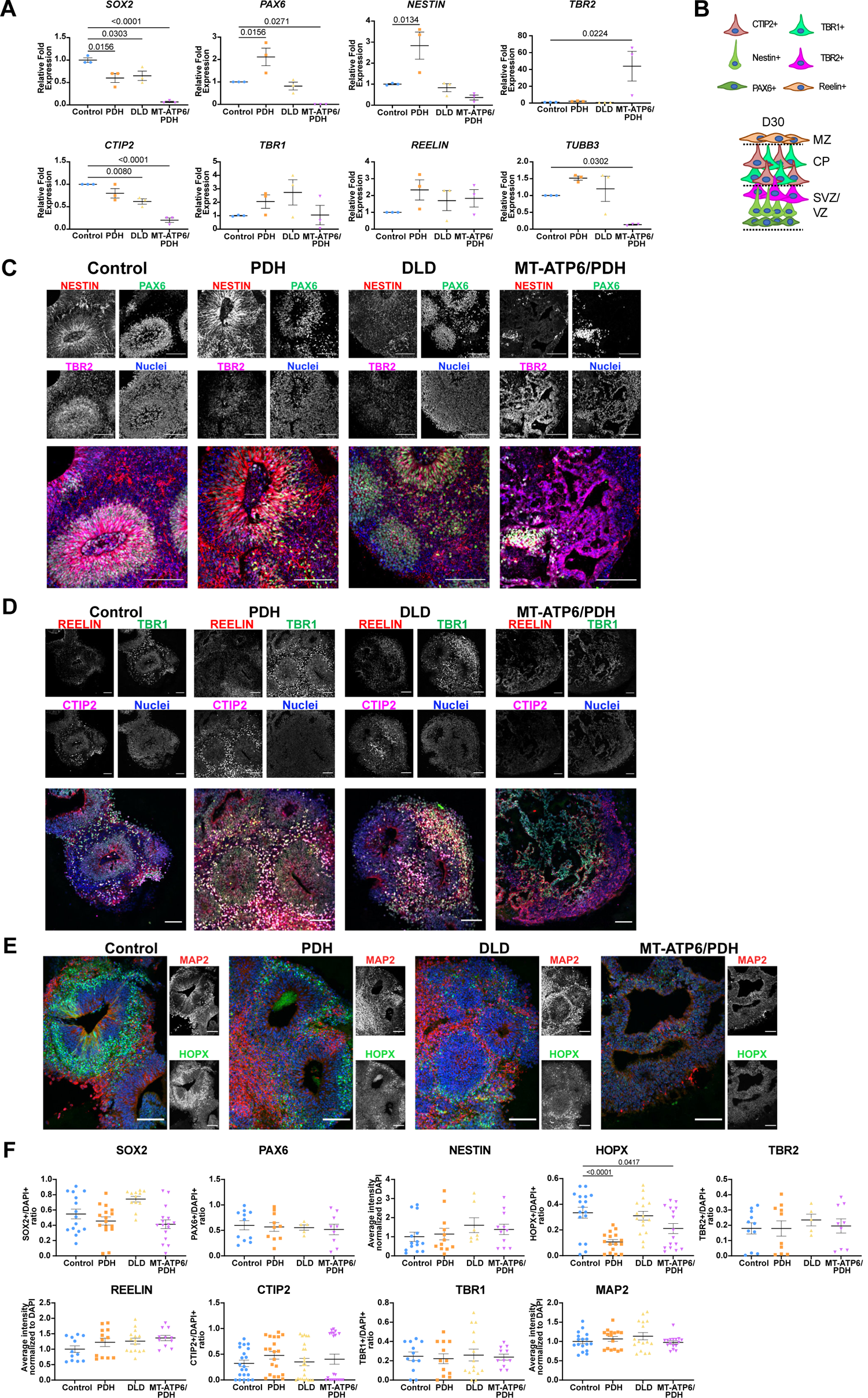
Leigh syndrome brain organoids show defects in SVZ/VZ and CP formation. A. qPCR quantification of the day 30 organoids. Neural progenitor cell populations were evaluated by the expression of *SOX2*, *PAX6* and *NESTIN*. Intermediate progenitor cells were identified with the marker *TBR2*. Marginal zone marker *REELIN*, cortical plate markers *TBR1* and *CTIP2*, and neuronal marker *TUBB3* were also evaluated. Fold change normalized to GPI and GAPDH as house-keeping genes. B. Schematic representation of the expected organization of the brain organoids on day 30. C-E. Representative immunostaining confocal images of day 30 brain organoids. MT-ATP6/PDH mutant presents severe disorganization of the SVZ/VZ markers PAX6 and TBR2, as well as the neural progenitor marker NESTIN (C). Cajal-Retzius neurons positive for REELIN were observed in the surface of the organoids (D). Cortical plate markers CTIP2 and TBR1 (D), as well as outer radial glia marker HOPX and the neuronal marker MAP2 (E). For E, nuclei in merge image correspond to the blue channel. Scale bar: 100μm. Images were generated from at least three different organoids per genotype from 3 independent organoid batches. F. Quantification of day immunofluorescence staining for day 30 brain organoids. Outer radial glia marker HOPX was reduced PDH mutant (p<0.0001) and MT-ATP6/PDH mutant (p=0.0417) derived brain organoids. SVZ: subventricular zone, VZ: ventricular zone, CP: cortical plate, MZ: marginal zone

**Supplemental Figure 6, Related to Figure 5.**
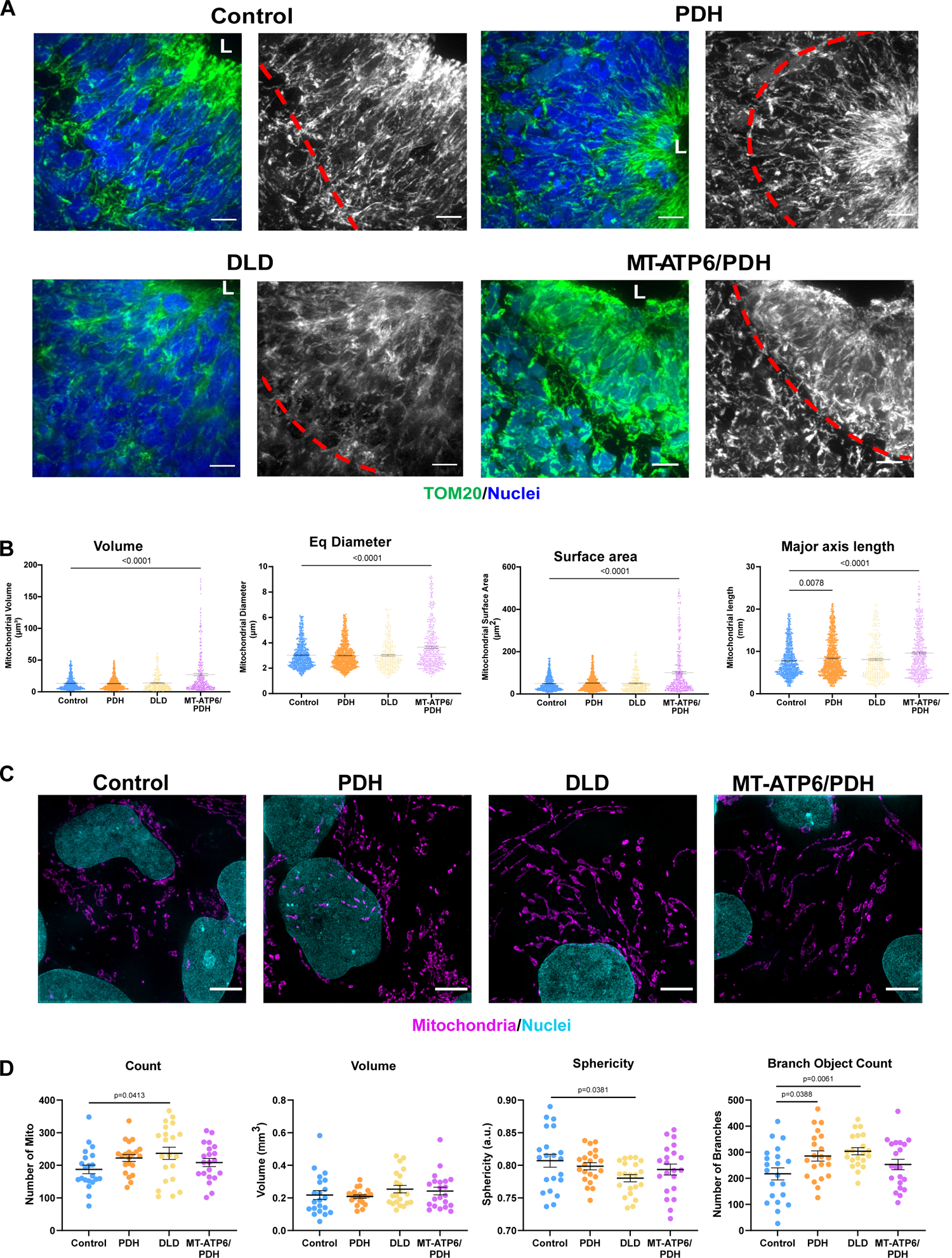
Leigh syndrome derived organoids show defects in mitochondrial morphology in the SVZ compartment. A. Representative confocal images of day 30 brain organoids showing mitochondrial morphology (TOM20). The red line divides the Sox2+ neural progenitor cells surrounding the lumen (L) from newly committed neurons. MT-ATP6/PDH mutant organoids show disorganization and fragmentation of the mitochondrial network compared to control. Scale bar: 10μm. B. Quantification of average mitochondrial volume, diameter, surface area, and major axis length are shown. Graphs represent mean ± SEM from at least three independent SVZs per phenotype from 3 independent organoid batches. Quantification was performed by 3D reconstruction of the mitochondrial network in interest. C. Representative super-resolution images of mitochondrial morphology (human anti-mitochondria) in LS and control NPCs. Scale bar: 5μm. D. Quantification of average mitochondrial number, volume, mitochondrial sphericity, and mitochondrial branching are shown, in which a spherical object would have a value of 1.0. Graphs represent mean ± SEM from at least three independent experiments (n > 20 cells per genotype).

**Supplemental Figure 7, Related to Figure 7 and Supplemental Table 2.**
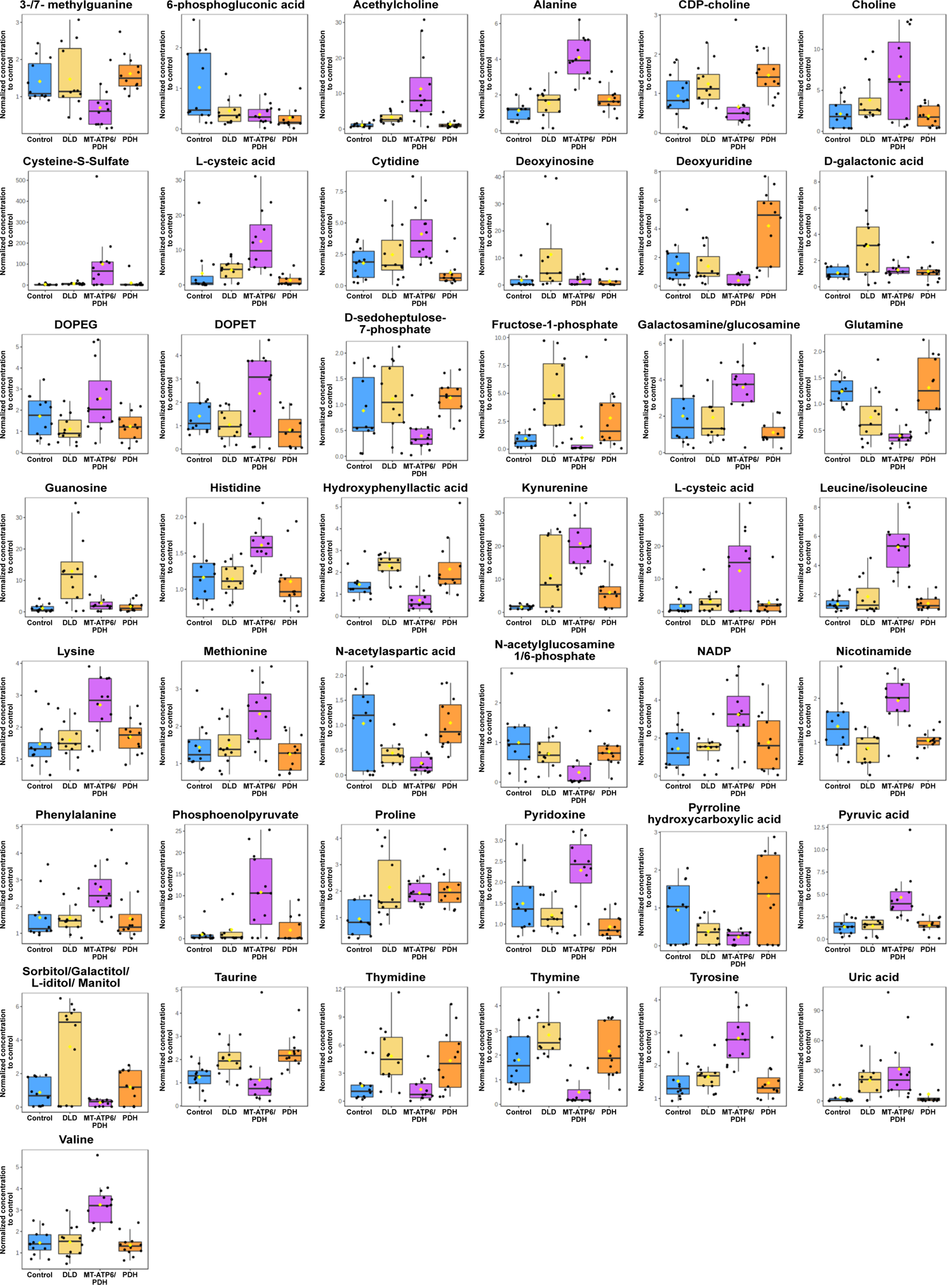
Day 40 LS organoids show changes in their metabolic profiles. Individual graph bars for the 43 metabolites identified as statistically dysregulated (p<0.05 and FDR of 0.01) in LS Organoids when compared to control. Statistical values can be found in Supplemental Table 2. A total of three batches of 40-day organoids per line (4 independent organoids per line per batch) were analyzed as described in methods.

**Supplemental Table 1, Related to Figure 1.** Summary characteristics of the Leigh syndrome patient derived fibroblast cell lines including the patient phenotype at diagnosis, the mutations identified, and published literature using the cell line.

| Cell line | Phenotype | Mutation | References |
| --- | --- | --- | --- |
| GM03672 | 1 y/o female, Caucasian.<br>Developmental regression; microcephalic;<br>elevated blood lactic acid and pyruvate;<br>only affected family member | PDH (E1) LOF<br>Pyruvate dehydrogenase<br>(79delC, Arg27fs) | Hinman et al. 1989;<br>Huh et al. 1990;<br>Johnson et al. 2019 |
| GM01503 | 3y/o female, Caucasian.<br>Lactic acidosis, psychomotor delay;<br>sister also affected.<br>Subnormal activation of pyruvate<br>dehydrogenase complex in<br>disrupted fibroblasts | DLD (E3)<br>Dihydrolipoamide<br>dehydrogenase<br>(A100G, Thr34Ala) | Cooper et al, 1969;<br>Murphy JV, 1973;<br>Sorbi& Blass, 1982;<br>Hinman et al. 1989;<br>Huh et al. 1990;<br>Vo et al. 2007;<br>Johnson et al. 2019 |
| GM13411 | 8 month old male, Chinese.<br>Lactic acidosis; developmental delay.<br>Hypertrophic cardiomyopathy.<br>Symmetric neural necrosis.<br>Fibroblast: 90% heteroplasmy | MT-ATP6<br>(T8993G, Leu156Arg)<br>PDH (E1) LOF<br>Pyruvate dehydrogenase<br>(79delC, Arg27fs) | Pastores et al. 1994;<br>Iyer et al.2012;<br>Ma et al. 2014;<br>Galera-Monge et al. 2016;<br>Zheng et al. 2016;<br>Johnson et al. 2019 |

**Supplemental Table 2, Related to Figure 7 and Supplemental Figure 7.**
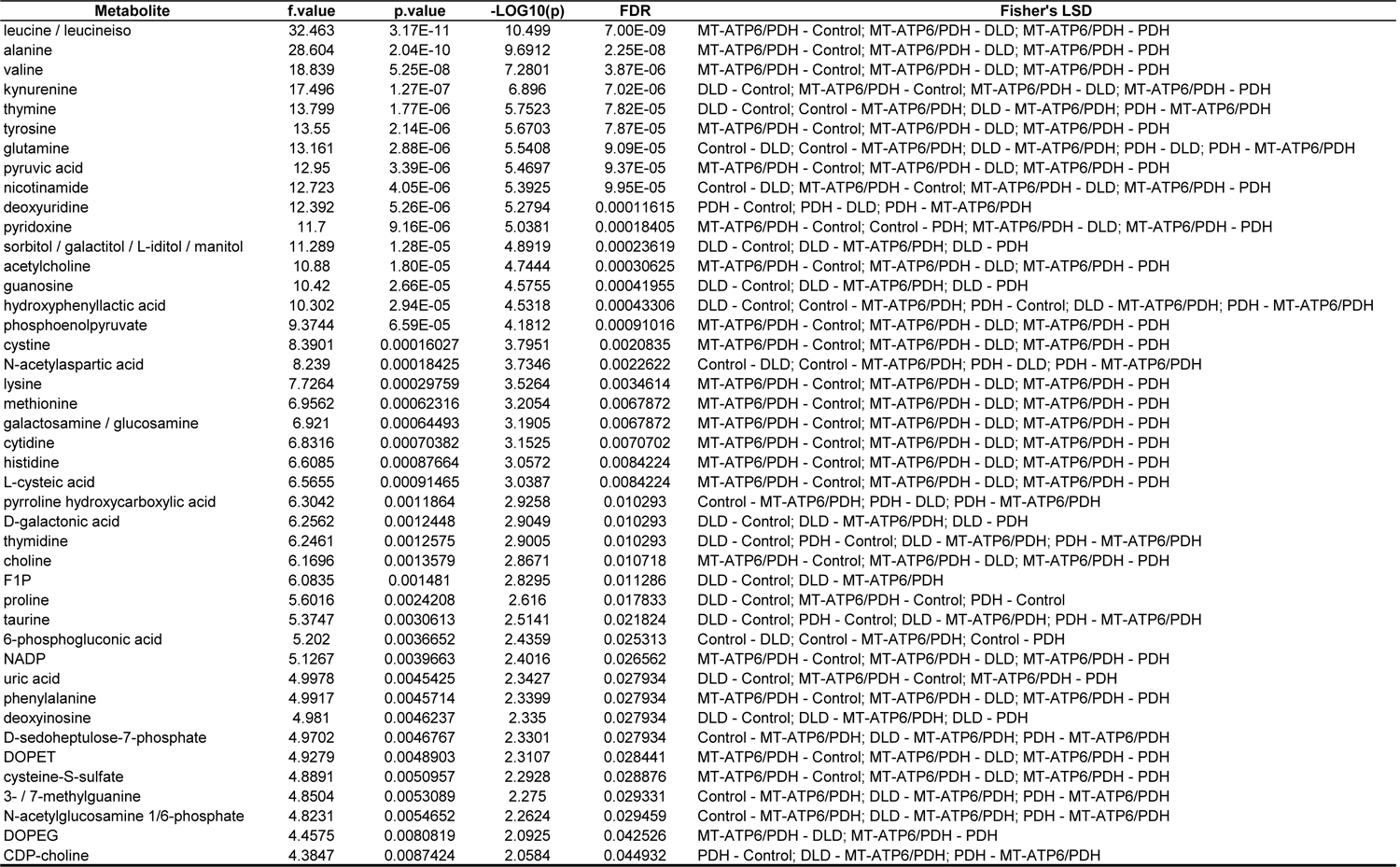
Dysregulated metabolites in day 40 LS derived cerebral organoids. LC-MS measured metabolite peak areas were normalized to the total ion count (TIC) by sample and fold change was determined by dividing each LS TIC normalized peak area by the control TIC normalized peak area for each metabolite. One way ANOVA was utilized to identified metabolites that were significantly dysregulated (p<0.05). Post-hoc comparison column using Fisher’s least significant difference method (Fisher’s LSD) shows the comparisons between different levels that are significant given the p value threshold. Results shown are averages for 3 independent runs with 4 individual organoids per phenotype per run. FDR: False Discovery Rate.

| Metabolite | f.value | p.value | -LOG10(p) | FDR | Fisher's LSD |
| --- | --- | --- | --- | --- | --- |
| leucine / leucineiso | 32.463 | 3.17E-11 | 10.499 | 7.00E-09 | MT-ATP6/PDH - Control; MT-ATP6/PDH - DLD; MT-ATP6/PDH - PDH |
| alanine | 28.604 | 2.04E-10 | 9.6912 | 2.25E-08 | MT-ATP6/PDH - Control; MT-ATP6/PDH - DLD; MT-ATP6/PDH - PDH |
| valine | 18.839 | 5.25E-08 | 7.2801 | 3.87E-06 | MT-ATP6/PDH - Control; MT-ATP6/PDH - DLD; MT-ATP6/PDH - PDH |
| kynurenine | 17.496 | 1.27E-07 | 6.896 | 7.02E-06 | DLD - Control; MT-ATP6/PDH - Control; MT-ATP6/PDH - DLD; MT-ATP6/PDH - PDH |
| thymine | 13.799 | 1.77E-06 | 5.7523 | 7.82E-05 | DLD - Control; Control - MT-ATP6/PDH; DLD - MT-ATP6/PDH; PDH - MT-ATP6/PDH |
| tyrosine | 13.55 | 2.14E-06 | 5.6703 | 7.87E-05 | MT-ATP6/PDH - Control; MT-ATP6/PDH - DLD; MT-ATP6/PDH - PDH |
| glutamine | 13.161 | 2.88E-06 | 5.5408 | 9.09E-05 | Control - DLD; Control - MT-ATP6/PDH; DLD - MT-ATP6/PDH; PDH - DLD; PDH - MT-ATP6/PDH |
| pyruvic acid | 12.95 | 3.39E-06 | 5.4697 | 9.37E-05 | MT-ATP6/PDH - Control; MT-ATP6/PDH - DLD; MT-ATP6/PDH - PDH |
| nicotinamide | 12.723 | 4.05E-06 | 5.3925 | 9.95E-05 | Control - DLD; MT-ATP6/PDH - Control; MT-ATP6/PDH - DLD; MT-ATP6/PDH - PDH |
| deoxyuridine | 12.392 | 5.26E-06 | 5.2794 | 0.00011615 | PDH - Control; PDH - DLD; PDH - MT-ATP6/PDH |
| pyridoxine | 11.7 | 9.16E-06 | 5.0381 | 0.00018405 | MT-ATP6/PDH - Control; Control - PDH; MT-ATP6/PDH - DLD; MT-ATP6/PDH - PDH |
| sorbitol / galactitol / L-iditol / manitol | 11.289 | 1.28E-05 | 4.8919 | 0.00023619 | DLD - Control; DLD - MT-ATP6/PDH; DLD - PDH |
| acetylcholine | 10.88 | 1.80E-05 | 4.7444 | 0.00030625 | MT-ATP6/PDH - Control; MT-ATP6/PDH - DLD; MT-ATP6/PDH - PDH |
| guanosine | 10.42 | 2.66E-05 | 4.5755 | 0.00041955 | DLD - Control; DLD - MT-ATP6/PDH; DLD - PDH |
| hydroxyphenyllactic acid | 10.302 | 2.94E-05 | 4.5318 | 0.00043306 | DLD - Control; Control - MT-ATP6/PDH; PDH - Control; DLD - MT-ATP6/PDH; PDH - MT-ATP6/PDH |
| phosphoenolpyruvate | 9.3744 | 6.59E-05 | 4.1812 | 0.00091016 | MT-ATP6/PDH - Control; MT-ATP6/PDH - DLD; MT-ATP6/PDH - PDH |
| cystine | 8.3901 | 0.00016027 | 3.7951 | 0.0020835 | MT-ATP6/PDH - Control; MT-ATP6/PDH - DLD; MT-ATP6/PDH - PDH |
| N-acetylaspartic acid | 8.239 | 0.00018425 | 3.7346 | 0.0022622 | Control - DLD; Control - MT-ATP6/PDH; PDH - DLD; PDH - MT-ATP6/PDH |
| lysine | 7.7264 | 0.00029759 | 3.5264 | 0.0034614 | MT-ATP6/PDH - Control; MT-ATP6/PDH - DLD; MT-ATP6/PDH - PDH |
| methionine | 6.9562 | 0.00062316 | 3.2054 | 0.0067872 | MT-ATP6/PDH - Control; MT-ATP6/PDH - DLD; MT-ATP6/PDH - PDH |
| galactosamine / glucosamine | 6.921 | 0.00064493 | 3.1905 | 0.0067872 | MT-ATP6/PDH - Control; MT-ATP6/PDH - DLD; MT-ATP6/PDH - PDH |
| cytidine | 6.8316 | 0.00070382 | 3.1525 | 0.0070702 | MT-ATP6/PDH - Control; MT-ATP6/PDH - DLD; MT-ATP6/PDH - PDH |
| histidine | 6.6085 | 0.00087664 | 3.0572 | 0.0084224 | MT-ATP6/PDH - Control; MT-ATP6/PDH - DLD; MT-ATP6/PDH - PDH |
| L-cysteic acid | 6.5655 | 0.00091465 | 3.0387 | 0.0084224 | MT-ATP6/PDH - Control; MT-ATP6/PDH - DLD; MT-ATP6/PDH - PDH |
| pyrroline hydroxycarboxylic acid | 6.3042 | 0.0011864 | 2.9258 | 0.010293 | Control - MT-ATP6/PDH; PDH - DLD; PDH - MT-ATP6/PDH |
| D-galactonic acid | 6.2562 | 0.0012448 | 2.9049 | 0.010293 | DLD - Control; DLD - MT-ATP6/PDH; DLD - PDH |
| thymidine | 6.2461 | 0.0012575 | 2.9005 | 0.010293 | DLD - Control; PDH - Control; DLD - MT-ATP6/PDH; PDH - MT-ATP6/PDH |
| choline | 6.1696 | 0.0013579 | 2.8671 | 0.010718 | MT-ATP6/PDH - Control; MT-ATP6/PDH - DLD; MT-ATP6/PDH - PDH |
| F1P | 6.0835 | 0.001481 | 2.8295 | 0.011286 | DLD - Control; DLD - MT-ATP6/PDH |
| proline | 5.6016 | 0.0024208 | 2.616 | 0.017833 | DLD - Control; MT-ATP6/PDH - Control; PDH - Control |
| taurine | 5.3747 | 0.0030613 | 2.5141 | 0.021824 | DLD - Control; PDH - Control; DLD - MT-ATP6/PDH; PDH - MT-ATP6/PDH |
| 6-phosphogluconic acid | 5.202 | 0.0036652 | 2.4359 | 0.025313 | Control - DLD; Control - MT-ATP6/PDH; Control - PDH |
| NADP | 5.1267 | 0.0039663 | 2.4016 | 0.026562 | MT-ATP6/PDH - Control; MT-ATP6/PDH - DLD; MT-ATP6/PDH - PDH |
| uric acid | 4.9978 | 0.0045425 | 2.3427 | 0.027934 | DLD - Control; MT-ATP6/PDH - Control; MT-ATP6/PDH - PDH |
| phenylalanine | 4.9917 | 0.0045714 | 2.3399 | 0.027934 | MT-ATP6/PDH - Control; MT-ATP6/PDH - DLD; MT-ATP6/PDH - PDH |
| deoxyinosine | 4.981 | 0.0046237 | 2.335 | 0.027934 | DLD - Control; DLD - MT-ATP6/PDH; DLD - PDH |
| D-sedoheptulose-7-phosphate | 4.9702 | 0.0046767 | 2.3301 | 0.027934 | Control - MT-ATP6/PDH; DLD - MT-ATP6/PDH; PDH - MT-ATP6/PDH |
| DOPET | 4.9279 | 0.0048903 | 2.3107 | 0.028441 | MT-ATP6/PDH - Control; MT-ATP6/PDH - DLD; MT-ATP6/PDH - PDH |
| cysteine-S-sulfate | 4.8891 | 0.0050957 | 2.2928 | 0.028876 | MT-ATP6/PDH - Control; MT-ATP6/PDH - DLD; MT-ATP6/PDH - PDH |
| 3- / 7-methylguanine | 4.8504 | 0.0053089 | 2.275 | 0.029331 | Control - MT-ATP6/PDH; DLD - MT-ATP6/PDH; PDH - MT-ATP6/PDH |
| N-acetylglucosamine 1/6-phosphate | 4.8231 | 0.0054652 | 2.2624 | 0.029459 | Control - MT-ATP6/PDH; DLD - MT-ATP6/PDH; PDH - MT-ATP6/PDH |
| DOPEG | 4.4575 | 0.0080819 | 2.0925 | 0.042526 | MT-ATP6/PDH - DLD; MT-ATP6/PDH - PDH |
| CDP-choline | 4.3847 | 0.0087424 | 2.0584 | 0.044932 | PDH - Control; DLD - MT-ATP6/PDH; PDH - MT-ATP6/PDH |

**Supplemental Table 3, Related to Related to Figure 7 and Supplemental Figure 7.**
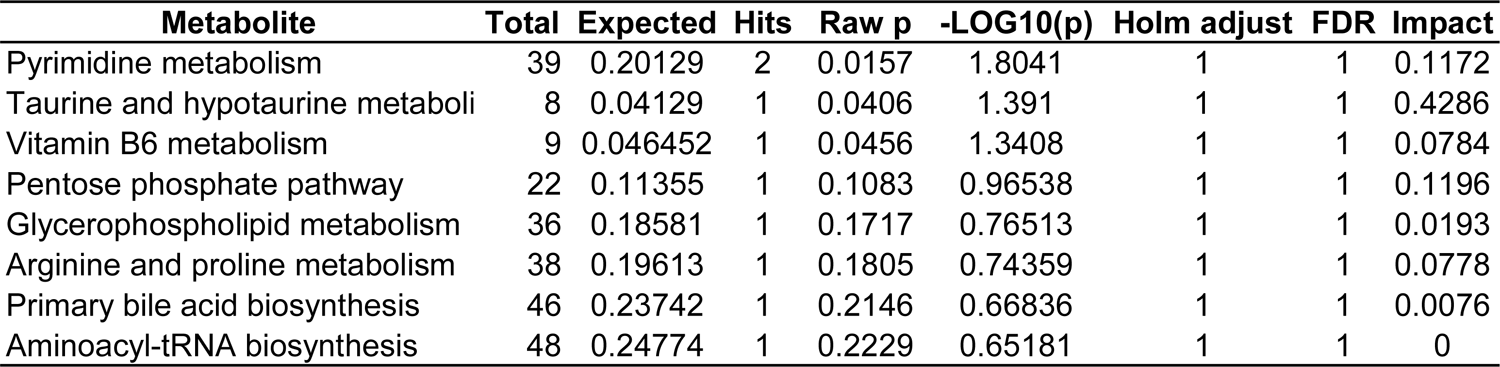
Summary of the metabolic pathways analysis for metabolites enriched in day 40 PDH brain organoids. Statistical p values from enrichment analysis are adjusted for multiple hypothesis testing. Total: total number of compounds in the pathway. Hits: matched number from the uploaded data. Raw p: original p value calculated from the enrichment analysis. Holm p: p value adjusted by Holm-Bonferroni method. FDR p: adjusted p value using False Discovery Rate. Impact: pathway impact value calculated from pathway topology analysis.

| **Metabolite** | **Total** | **Expected** | **Hits** | **Raw p** | **-LOG10(p)** | **Holm adjust** | **FDR** | **Impact** |
| --- | --- | --- | --- | --- | --- | --- | --- | --- |
| Pyrimidine metabolism | 39 | 0.20129 | 2 | 0.0157 | 1.8041 | 1 | 1 | 0.1172 |
| Taurine and hypotaurine metaboli | 8 | 0.04129 | 1 | 0.0406 | 1.391 | 1 | 1 | 0.4286 |
| Vitamin B6 metabolism | 9 | 0.046452 | 1 | 0.0456 | 1.3408 | 1 | 1 | 0.0784 |
| Pentose phosphate pathway | 22 | 0.11355 | 1 | 0.1083 | 0.96538 | 1 | 1 | 0.1196 |
| Glycerophospholipid metabolism | 36 | 0.18581 | 1 | 0.1717 | 0.76513 | 1 | 1 | 0.0193 |
| Arginine and proline metabolism | 38 | 0.19613 | 1 | 0.1805 | 0.74359 | 1 | 1 | 0.0778 |
| Primary bile acid biosynthesis | 46 | 0.23742 | 1 | 0.2146 | 0.66836 | 1 | 1 | 0.0076 |
| Aminoacyl-tRNA biosynthesis | 48 | 0.24774 | 1 | 0.2229 | 0.65181 | 1 | 1 | 0 |

**Supplemental Table 4, Related to Related to Figure 7 and Supplemental Figure 7.**
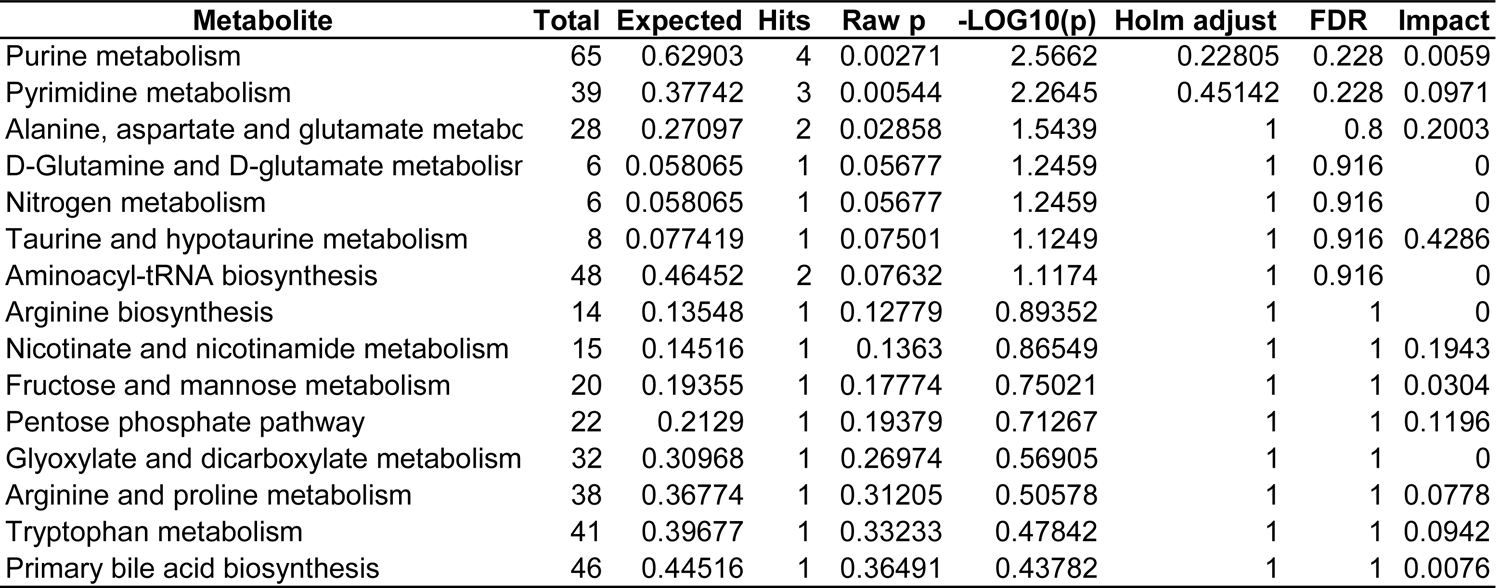
Summary of the metabolic pathways analysis for metabolites enriched in day 40 DLD brain organoids. Statistical p values from enrichment analysis are adjusted for multiple hypothesis testing. Total: total number of compounds in the pathway. Hits: matched number from the uploaded data. Raw p: original p value calculated from the enrichment analysis. Holm p: p value adjusted by Holm-Bonferroni method. FDR p: adjusted p value using False Discovery Rate. Impact: pathway impact value calculated from pathway topology analysis.

| Metabolite | Total | Expected | Hits | Raw p | -LOG10(p) | Holm adjust | FDR | Impact |
| --- | --- | --- | --- | --- | --- | --- | --- | --- |
| Purine metabolism | 65 | 0.62903 | 4 | 0.00271 | 2.5662 | 0.22805 | 0.228 | 0.0059 |
| Pyrimidine metabolism | 39 | 0.37742 | 3 | 0.00544 | 2.2645 | 0.45142 | 0.228 | 0.0971 |
| Alanine, aspartate and glutamate metabo | 28 | 0.27097 | 2 | 0.02858 | 1.5439 | 1 | 0.8 | 0.2003 |
| D-Glutamine and D-glutamate metabolisr | 6 | 0.058065 | 1 | 0.05677 | 1.2459 | 1 | 0.916 | 0 |
| Nitrogen metabolism | 6 | 0.058065 | 1 | 0.05677 | 1.2459 | 1 | 0.916 | 0 |
| Taurine and hypotaurine metabolism | 8 | 0.077419 | 1 | 0.07501 | 1.1249 | 1 | 0.916 | 0.4286 |
| Aminoacyl-tRNA biosynthesis | 48 | 0.46452 | 2 | 0.07632 | 1.1174 | 1 | 0.916 | 0 |
| Arginine biosynthesis | 14 | 0.13548 | 1 | 0.12779 | 0.89352 | 1 | 1 | 0 |
| Nicotinate and nicotinamide metabolism | 15 | 0.14516 | 1 | 0.1363 | 0.86549 | 1 | 1 | 0.1943 |
| Fructose and mannose metabolism | 20 | 0.19355 | 1 | 0.17774 | 0.75021 | 1 | 1 | 0.0304 |
| Pentose phosphate pathway | 22 | 0.2129 | 1 | 0.19379 | 0.71267 | 1 | 1 | 0.1196 |
| Glyoxylate and dicarboxylate metabolism | 32 | 0.30968 | 1 | 0.26974 | 0.56905 | 1 | 1 | 0 |
| Arginine and proline metabolism | 38 | 0.36774 | 1 | 0.31205 | 0.50578 | 1 | 1 | 0.0778 |
| Tryptophan metabolism | 41 | 0.39677 | 1 | 0.33233 | 0.47842 | 1 | 1 | 0.0942 |
| Primary bile acid biosynthesis | 46 | 0.44516 | 1 | 0.36491 | 0.43782 | 1 | 1 | 0.0076 |

**Supplemental Table 5, Related to Related to Figure 7 and Supplemental Figure 7.** Summary of the metabolic pathways analysis for metabolites enriched in day 40 MT-ATP6/PDH brain organoids. Statistical p values from enrichment analysis are adjusted for multiple hypothesis testing. Total: total number of compounds in the pathway. Hits: matched number from the uploaded data. Raw p: original p value calculated from the enrichment analysis. Holm p: p value adjusted by Holm-Bonferroni method. FDR p: adjusted p value using False Discovery Rate. Impact: pathway impact value calculated from pathway topology analysis.

| Metabolite | Total | Expected | Hits | Raw p | -LOG10(p) | Holm adjust | FDR | Impact |
| --- | --- | --- | --- | --- | --- | --- | --- | --- |
| Aminoacyl-tRNA biosynthesis | 48 | 0.71226 | 6 | 4.35E-05 | 4.3617 | 0.0036523 | 0.0036523 | 0 |
| Alanine, aspartate and glutamate metabolism | 28 | 0.41548 | 4 | 0.00059684 | 3.2241 | 0.049538 | 0.025067 | 0.20032 |
| Cysteine and methionine metabolism | 33 | 0.48968 | 3 | 0.011653 | 1.9336 | 0.95556 | 0.32629 | 0.10446 |
| Arginine and proline metabolism | 38 | 0.56387 | 3 | 0.017166 | 1.7653 | 1 | 0.33051 | 0.13058 |
| Nicotinate and nicotinamide metabolism | 15 | 0.22258 | 2 | 0.019673 | 1.7061 | 1 | 0.33051 | 0.1943 |
| Citrate cycle (TCA cycle) | 20 | 0.29677 | 2 | 0.034034 | 1.4681 | 1 | 0.47648 | 0.04634 |
| Pyruvate metabolism | 22 | 0.32645 | 2 | 0.040643 | 1.391 | 1 | 0.48771 | 0.20684 |
| Glycolysis / Gluconeogenesis | 26 | 0.38581 | 2 | 0.055171 | 1.2583 | 1 | 0.54228 | 0.20594 |
| Phenylalanine, tyrosine and tryptophan biosynthesis | 4 | 0.059355 | 1 | 0.058102 | 1.2358 | 1 | 0.54228 | 0.5 |
| Glyoxylate and dicarboxylate metabolism | 32 | 0.47484 | 2 | 0.079815 | 1.0979 | 1 | 0.55522 | 0 |
| Glycine, serine and threonine metabolism | 33 | 0.48968 | 2 | 0.084213 | 1.0746 | 1 | 0.55522 | 0 |
| D-Glutamine and D-glutamate metabolism | 6 | 0.089032 | 1 | 0.085928 | 1.0659 | 1 | 0.55522 | 0 |
| Nitrogen metabolism | 6 | 0.089032 | 1 | 0.085928 | 1.0659 | 1 | 0.55522 | 0 |
| Glycerophospholipid metabolism | 36 | 0.53419 | 2 | 0.097841 | 1.0095 | 1 | 0.5848 | 0.02582 |
| Pyrimidine metabolism | 39 | 0.57871 | 2 | 0.11206 | 0.95053 | 1 | 0.5848 | 0.03727 |
| Taurine and hypotaurine metabolism | 8 | 0.11871 | 1 | 0.11297 | 0.94705 | 1 | 0.5848 | 0 |
| Ubiquinone and other terpenoid-quinone biosynthesis | 9 | 0.13355 | 1 | 0.1262 | 0.89895 | 1 | 0.5848 | 0 |
| Tyrosine metabolism | 42 | 0.62323 | 2 | 0.12681 | 0.89685 | 1 | 0.5848 | 0.13972 |
| Phenylalanine metabolism | 10 | 0.14839 | 1 | 0.13924 | 0.85624 | 1 | 0.5848 | 0 |
| Biotin metabolism | 10 | 0.14839 | 1 | 0.13924 | 0.85624 | 1 | 0.5848 | 0 |
| Arginine biosynthesis | 14 | 0.20774 | 1 | 0.18957 | 0.72223 | 1 | 0.75827 | 0 |
| Purine metabolism | 65 | 0.96452 | 2 | 0.25062 | 0.60098 | 1 | 0.9489 | 0 |
| Selenocompound metabolism | 20 | 0.29677 | 1 | 0.25982 | 0.58533 | 1 | 0.9489 | 0 |
| Pentose phosphate pathway | 22 | 0.32645 | 1 | 0.28191 | 0.54989 | 1 | 0.98669 | 0.11955 |
| Lysine degradation | 25 | 0.37097 | 1 | 0.31387 | 0.50324 | 1 | 1 | 0 |
| Glutathione metabolism | 28 | 0.41548 | 1 | 0.34447 | 0.46285 | 1 | 1 | 0.0018 |
| Tryptophan metabolism | 41 | 0.60839 | 1 | 0.46262 | 0.33477 | 1 | 1 | 0.09417 |

## Notes

### Competing Interest Statement

The authors have declared no competing interest.

